# A Probabilistic, Distributed, Recursive Mechanism for Decision-making in the Brain

**DOI:** 10.1101/036277

**Authors:** Javier A. Caballero, Mark D. Humphries, Kevin N. Gurney

## Abstract

Decision formation recruits many brain regions, but the procedure they jointly execute is unknown. Here we characterize its essential composition, using as a framework a novel recursive Bayesian algorithm that makes decisions based on spike-trains with the statistics of those in sensory cortex (MT). Using it to simulate the random-dot-motion task, we demonstrate it quantitatively replicates the choice behaviour of monkeys, whilst predicting losses of otherwise usable information from MT. Its architecture maps to the recurrent cortico-basal-ganglia-thalamo-cortical loops, whose components are all implicated in decision-making. We show that the dynamics of its mapped computations match those of neural activity in the sensorimotor cortex and striatum during decisions, and forecast those of basal ganglia output and thalamus. This also predicts which aspects of neural dynamics are and are not part of inference. Our single-equation algorithm is probabilistic, distributed, recursive, and parallel. Its success at capturing anatomy, behaviour, and electrophysiology suggests that the mechanism implemented by the brain has these same characteristics.

**Author Summary:** Decision-making is central to cognition. Abnormally-formed decisions characterize disorders like over-eating, Parkinson’s and Huntington’s diseases, OCD, addiction, and compulsive gambling. Yet, a unified account of decisionmaking has, hitherto, remained elusive. Here we show the essential composition of the brain’s decision mechanism by matching experimental data from monkeys making decisions, to the knowable function of a novel statistical inference algorithm. Our algorithm maps onto the large-scale architecture of decision circuits in the primate brain, replicating the monkeys’ choice behaviour and the dynamics of the neural activity that accompany it. Validated in this way, our algorithm establishes a basic framework for understanding the mechanistic ingredients of decisionmaking in the brain, and thereby, a basic platform for understanding how pathologies arise from abnormal function.

## Introduction

Decisions rely on evidence that is collected for, accumulated about, and contrasted between available options. Neural activity consistent with evidence accumulation over time has been reported in parietal and frontal sensorimotor cortex [1–5], and in the subcortical striatum [6,7]. What overall computation underlies these local snapshots, and how it is distributed across cortical and subcortical circuits, is unknown.

Multiple models of decision making match aspects of recorded choice behaviour, associated neural activity or both [8–16]. While successful, they lack insight into the underlying decision mechanism. In contrast, other studies have shown how exact inference algorithms may be plausibly implemented by a range of neural circuits [17–21]; however, none of these has reproduced experimental decision data.

Here we test the hypothesis that the brain implements an approximation to an exact inference algorithm for decision making. We show that the algorithm reproduces behaviour quantitatively while the dynamics of its inner variables match those of corresponding neural signals on the random dot motion task —a highly developed paradigm to probe decision formation. By doing so, we predict how experimentally-acquired snapshots of neural activity map onto inference operations. We show this mapping accounts for the involvement of full recurrent cortico-subcortical loops in decision making. Evidence accumulation is thus predicted to occur over the entire loops, not just within cortex. Introducing this algorithm enables us to predict which aspects of neural activity are necessary for inference —hence decision-making— and which are not. For instance, recent data questioned whether non-increasing cortical firing rates encode evidence accumulation during decisions [22,23]. We demonstrate that, counter-intuitively, nonincreasing as well as increasing cortical rates can encode likelihood functions, and hence evidence accumulation.

Our algorithm explains the decision-correlated experimental data more comprehensively than any prior model, thus introducing a new, cohesive formal framework to interpret it. Collectively, our analyses and simulations indicate that mammalian decision-making is implemented as a probabilistic, recursive, parallel procedure distributed across the cortico-basal-ganglia-thalamo-cortical loops.

## Results

We tested our algorithm against behavioural and electrophysiological data recorded in sensorimotor cortex [3] and striatum [6], from monkeys performing 2- and 4-alternative reaction-time versions of the random dot motion task (Fig 1b,c). The decision evidence for the algorithm also simulates spike-trains from sensory cortex (the area that provides evidence to sensorimotor cortex), whose statistics we extracted from a third random-dot-task data set by [24]. In all forms of the task, the monkey observes the motion of dots and indicates the dominant direction of motion with a saccadic eye movement to a target in that direction. Task difficulty is controlled by the coherence of the motion: the percentage of dots moving in the target’s direction.

**Figure 1:**
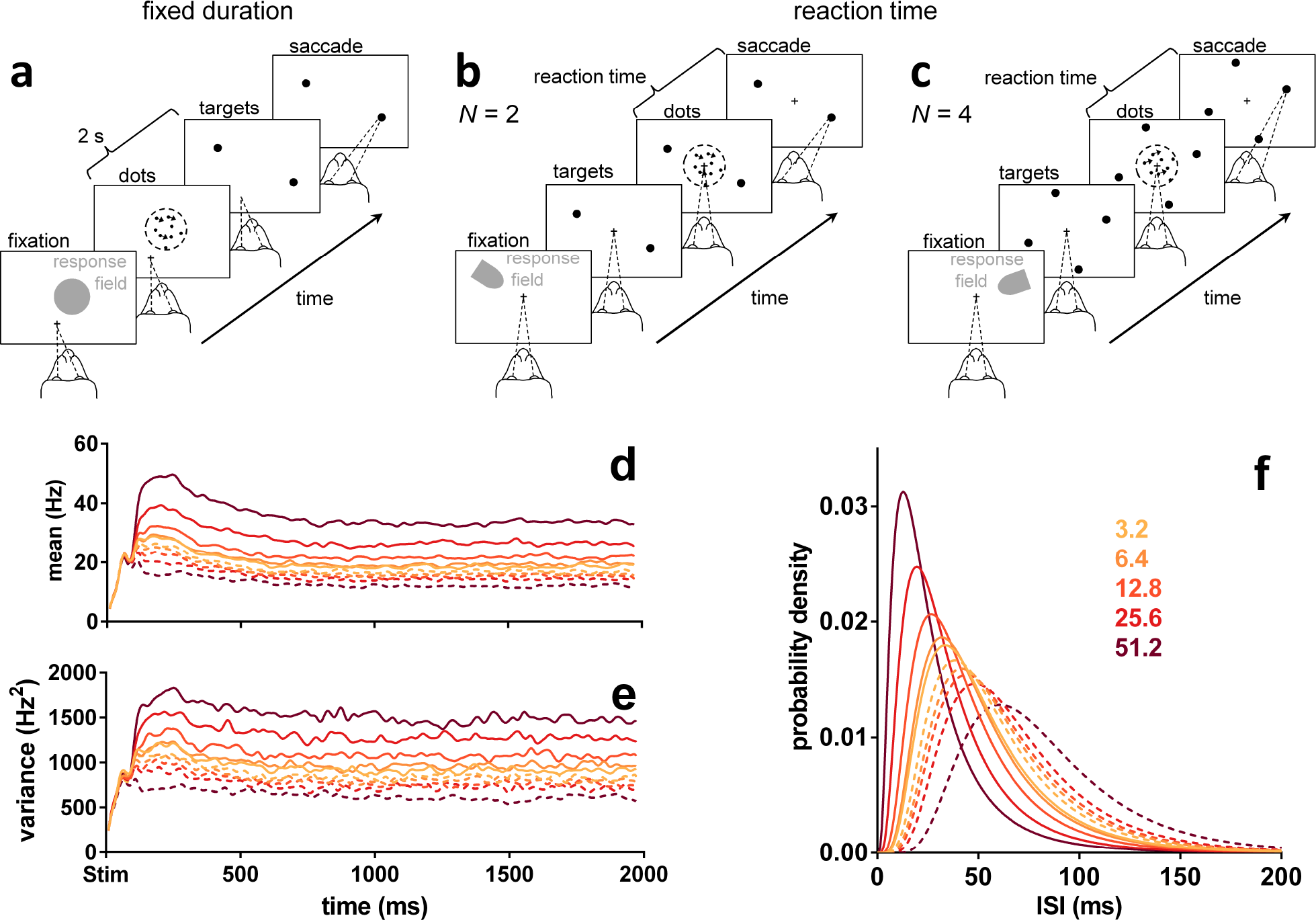
Random dot motion task and statistics of neural responses in sensory cortex (MT). (a) Fixed duration task for MT recordings [24]. (b, c) Reaction time task for sensorimotor cortex and striatum recordings, *N* = 2, 4 alternatives [3,6]. (d, e) Smoothed population moving mean and variance of the firing rate of MT during the fixed duration dot motion task (189–213 neurons), aligned at onset of the dot stimulus (Stim), for a variety of coherence percentages (colour-coded as in the legend in panel f). Solid lines are statistics when dots were moving in the preferred direction of the MT neuron. Dashed lines are statistics when dots were moving in the opposite, null direction. Data from [24], re-analysed. (f) Lognormal density functions for the inter-spike intervals (ISI) specified by the statistics over the approximately stationary segment of (d, e) before smoothing (parameter set Ω in Table 1). Preferred and null motion directions by line type as in (d, e).

During the dot motion task, neurons in the middle-temporal visual area (MT) respond more vigorously to visual stimuli moving in their “preferred” direction than in the opposite “null” direction [24]. Both the mean (Fig 1d) and variance (Fig 1e) of their response are proportional to the coherence of the motion (see also S4 Fig). MT responses are thence assumed to be the uncertain evidence upon which a choice is made in this task [1,9].

### Recursive MSPRT

Normative algorithms are useful benchmarks to test how well the brain approximates an optimal probabilistic computation. The family of the multi-hypothesis sequential probability ratio test (MSPRT) [25] is an attractive normative framework for understanding decision-making. However, the MSPRT is a feedforward algorithm. It cannot account for the ubiquitous presence of feedback in neural circuits and, as we show ahead, for slow dynamics in neural activity that result from this recurrence during decisions. To solve this, we introduce a novel recursive generalization, the rMSPRT, which uses a generalized, feedback form of the Bayes’ rule we deduced here from first principles (Eq 5).

We now conceptually review the MSPRT and introduce the rMSPRT (Fig 2), giving full mathematical definitions and deductions in the Materials and Methods. The (r)MSPRT decides which of N parallel, competing alternatives (or hypotheses) is the best choice, based on C sequentially sampled streams of evidence (or data). For modelling the dot-motion task, we have *N* = 2 or *N* = 4 hypotheses —the possible saccades to available targets (Fig 1b,c)— and the C uncertain *evidence streams* are assumed to be simultaneous spike-trains produced by visual-motion-sensitive MT neurons [1,9] (see Fig 9 in the Methods). Every time new evidence arrives, the (r)MSPRT refreshes ‘on-line’ the likelihood of each hypothesis: the plausibility of the combined evidence streams assuming that hypothesis is true. The likelihood is then multiplied by the probability of that hypothesis based on past experience (the prior). This product for every hypothesis is then normalized by the sum of the products from all N hypotheses; normalisation is crucial for decision, as it provides the competition between hypotheses. The result is the probability of each hypothesis given current evidence (the posterior) —a *decision variable* per hypothesis. Finally, posteriors are compared to a threshold, whose position controls the speed-accuracy trade-off. A decision is then made to either choose the most probable hypothesis, if its posterior surpassed the threshold, or to continue sampling the evidence streams otherwise. Crucially, the (r)MSPRT allows us to use the same algorithm irrespective of the number of alternatives, and thus aim at a unified explanation of the *N* = 2 and *N* = 4 dot-motion task variants.

**Figure 2:**
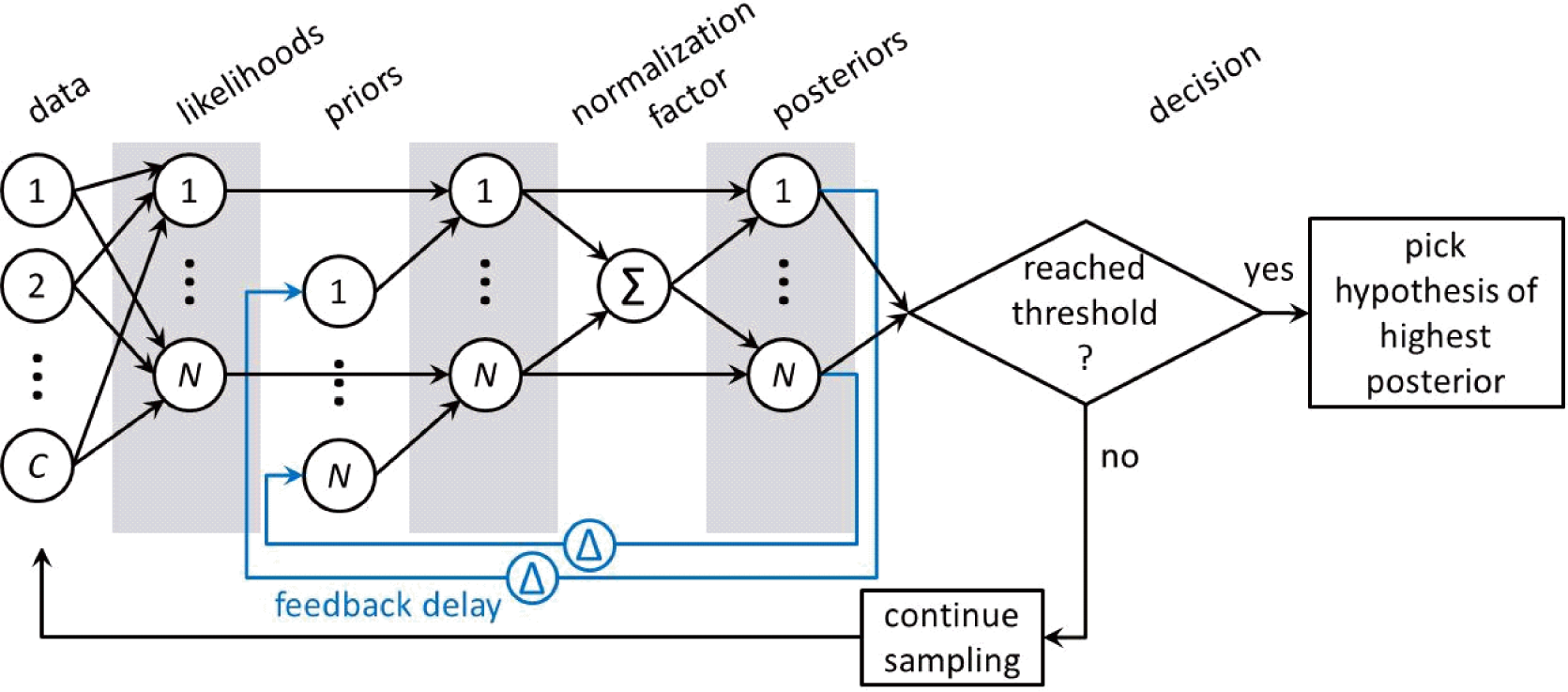
The MSPRT and rMSPRT as a diagram. Circles joined by arrows are the Bayes’ rule. All *C* evidence streams (data) are used to compute every one of the N likelihood functions. The product of the likelihood and prior probability of every hypothesis is normalized by the sum (Σ) of all products of likelihoods and priors, to produce the posterior probability of that hypothesis. All posteriors are then compared to a constant threshold. A decision is made every time with two possible outcomes: if a posterior reached the threshold, the hypothesis with the highest posterior is picked, otherwise, sampling from the evidence streams continues. The MSPRT as in [25] and [17] only requires what is shown in black. The general recursive MSPRT introduced here re-uses the posteriors Δ time steps in the past for present inference, thus re-using itself; hence the rMSPRT is shown in black and blue. If we are to work with the negative-logarithm of the Bayes’ rule —as we do in this article— all relations between computations are preserved, but products of computations become additions of their logarithms and the divisions for normalization become their negative logarithm. Eq 9 shows this for the rMSPRT. The rMSPRT itself is formalized by Eq 10.

The MSPRT is a special case of the rMSPRT (in its general form in Eqs 5 and 10) when priors do not change or, equivalently, for an infinite recursion delay; that is, Δ → ∞. Also, the previous recurrent extension of MSPRT [18,26] is a special case of the rMSPRT when Δ = 1. Hence, our rMSPRT generalizes both in allowing the re-use of posteriors from *any* given time in the past as priors for present inference. This uniquely allows us to map the rMSPRT onto neural circuits containing arbitrary feedback delays, in particular solving the problem of decomposing the decision-making algorithm into distributed components across multiple brain regions. We show below how this allows us to map the rMSPRT onto the cortico-basal-ganglia-thalamo-cortical loops.

Inference using recursive and non-recursive forms of Bayes’ rule gives the same results (*e.g.* see [27]), and so MSPRT and rMSPRT perform identically. Thus, like MSPRT [17,25], for *N* = 2 rMSPRT also collapses to the sequential probability ratio test of [28]; the rMSPRT is thereby optimal, not only in the oft-used sense of using all available information to do statistical inference (*e.g.* using the Bayes’ rule), but also in the strict sense that it requires the smallest expected number of observations, thus the shortest time to decide, at any given error rate (which follows from [29]). This is to say that the (r)MSPRT is quasi-Bayesian in general: the physical limit of performance or ideal Bayesian observer for two-alternative decisions (*N* = 2), and an asymptotic approximation to it for decisions between more than two (N > 2) (which follows from [17,25]).

### Upper bounds of decision time predicted by the (r)MSPRT

The hypothesis that the brain approximates an exact inference algorithm during decision formation is so far untested. This requires showing how uncertain sensory spike-trains can be transformed into the experimentally recorded choices. We do so here for the first time by comparing the predicted choice reaction times of the (r)MSPRT to those of monkeys performing the random dot motion task. We sought to account for the reaction time dependence on three factors: the coherence of the dot motion, the number of decision alternatives, and the trial’s outcome (error, correct). We use a particular instance of rMSPRT (Eqs 9 and 10) to determine predicted normative bounds on the decision time in the dot motion task. We can then ask how well monkeys approximate such bounds. The bounds result from using a minimal amount of sensory information, by assuming as many evidence streams (spike-trains from MT neurons) as alternatives; that is, C = N. Thus, this rMSPRT instance gives the upper bound on optimal expected decision times (exact for *N* = 2 alternatives, approximate for *N* = 4) per *condition* (given combination of coherence and N). Assuming *C* > *N* would predict even shorter optimal expected decision times (see [20]).

We assume that during the random dot motion task (Fig 1a-c), the evidence streams for every possible saccade come as simultaneous sequences of inter-spike intervals (ISI) produced in MT. On each time step, fresh evidence is drawn from the appropriate (null or preferred direction; see also Fig 9) ISI distributions extracted from MT data (Fig 1f). By repeating the simulations for thousands of trials per condition, we can compare algorithm and monkey performance.

Using these data-determined MT statistics, the (r)MSPRT predicts that the mean decision time on the dot motion task is a decreasing function of coherence (Fig 3a). For comparison with monkey reaction times, the algorithm’s reaction times are the sum of its decision times and estimated non-decision time, encompassing sensory delays and motor execution. For macaques 200–300 ms of non-decision time is a plausible range [30,31]. Within this range, monkeys tend not to reach the predicted upper bound of reaction time (Fig 3a).

**Figure 3:**
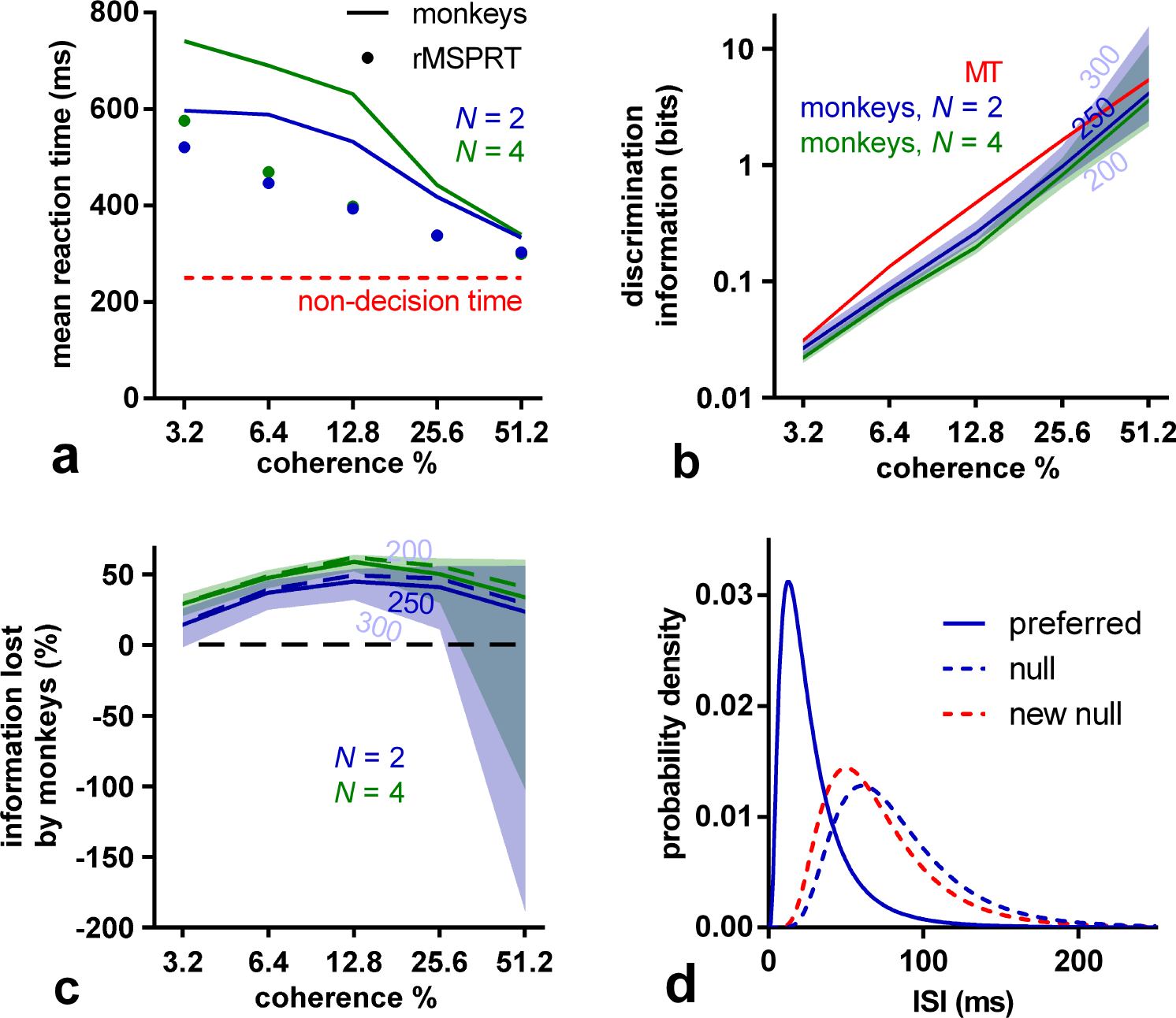
(r)MSPRT predicts information loss during decision making. (a) Comparison of the mean reaction time of monkeys for 2 and 4 alternatives (lines) with that predicted by (r)MSPRT (markers), both for correct trials. Red line: assumed 250 ms of non-decision time. Simulation values are means over 100 Monte Carlo experiments each comprising 3200, 4800 total trials for *N* = 2,4, correspondingly, under the parameter set Ω extracted from MT recordings. (b) Discrimination information per ISI in MT statistics (red) compared to the (r)MSPRT’s predictions of the discrimination information available to the monkeys (blue, green). Central lines are for a non-decision time of 250 ms; the edges of the correspondingly-coloured shaded regions are for non-decision times of 300 and 200 ms. (c) As per panel (b), but expressed as a percentage of information lost by monkeys with respect to the information available in MT for the three assumed non-decision times (solid lines and shadings). The information lost if the reaction time match is further enhanced is shown as dashed lines (assuming 250 ms of non-decision time; see Materials and Methods). (d) Example ISI density functions before (blue) and after (solid blue and dashed red) information depletion; *N* = 2, 51.2 % coherence, and 250 ms of non-decision time. The null distribution was adjusted to become the ‘new null’ by changing its mean and standard deviation to make it more similar to the preferred distribution. Once done throughout and for a non-decision time of 250 ms, this procedure gives ISI distributions bearing a reduced amount of discrimination information (blue or green lines in panel b), rather than the full discrimination information actually produced by MT (red line). That is, after adjustment, the discrimination information between the preferred and ‘new null’ distributions matches that estimated from the monkeys’ performance.

### The monkey brain loses otherwise useful information from sensory evidence

The (r)MSPRT framework suggests that decision times directly depend on the discrimination information in the evidence. Discrimination information here is measured as the divergence between pairs of distributions of ISIs (those in Fig 1f) produced simultaneously by MT neurons responding to the same stimulus: one where they were tuned to the dominant motion direction of the dots (it was their preferred; solid lines in Fig 1f) and another where they were not (it was a null direction; dashed lines). Intuitively, the larger this divergence or difference, the easier and hence faster the decision. We can estimate how much discrimination information monkeys used by asking how much the exact inference performed by (r)MSPRT would require to obtain the same reaction times on correct trials as the monkeys, per condition. We thus find, first, that the discrimination information available for decision is very similar across N (Fig 3b), implying that monkeys use MT sensory information consistently. Second, and most important, we find that monkeys tended to use less discrimination information than that in ISI distributions in their MT when making the decision. In contrast, the (r)MSPRT uses the full discrimination information available. This implies that the decision-making mechanism in the monkey brain lost large proportions of MT discrimination information (Fig 3c). Since these (r)MSPRT decision times are upper bounds, this in turn means that this loss of discrimination information in monkeys (Fig 3c) is the minimum.

### (r)MSPRT with depleted information quantitatively reproduces monkey performance

To verify if this information loss alone could account for the monkeys’ deviation from the (r)MSPRT upper bounds, we depleted the discrimination information of its input distributions to exactly match the estimated monkey loss in Fig 3c per condition. We did so only by modifying the mean and standard deviation of the null direction ISI distribution, to make it more similar to the preferred distribution (exemplified in Fig 3d).

Using these information-depleted statistics, the mean reaction times predicted by the (r)MSPRT in correct trials closely match those of monkeys (Fig 4a). Importantly, this involved no parameter fitting. Instead, we used the fact that for (r)MSPRT the mean total information for a decision is constant given error rate and *N*; this implies that longer decision times could only result from reducing the discrimination information in the evidence. Strikingly, although this information-depletion procedure is based only on data from correct trials, the (r)MSPRT now also matches closely the mean reaction times of monkeys from error trials (Fig 4b), which are consistently longer than those of correct trials (S1 Fig). Moreover, for both correct and error trials the (r)MSPRT accurately captures the relative scaling of mean reaction time by the number of alternatives (Fig 4a,b).

The reaction time distributions of the algorithm closely resemble those of monkeys in that they are positively skewed and exhibit shorter right tails for higher coherence levels (Fig 4c-f). These qualitative features are captured across both correct and error trials, and 2 and 4-alternative tasks. Together, these results support the hypothesis that the primate brain approximates an algorithm similar to the rMSPRT, ‘starved’ of sensory discrimination information.

**Figure 4:**
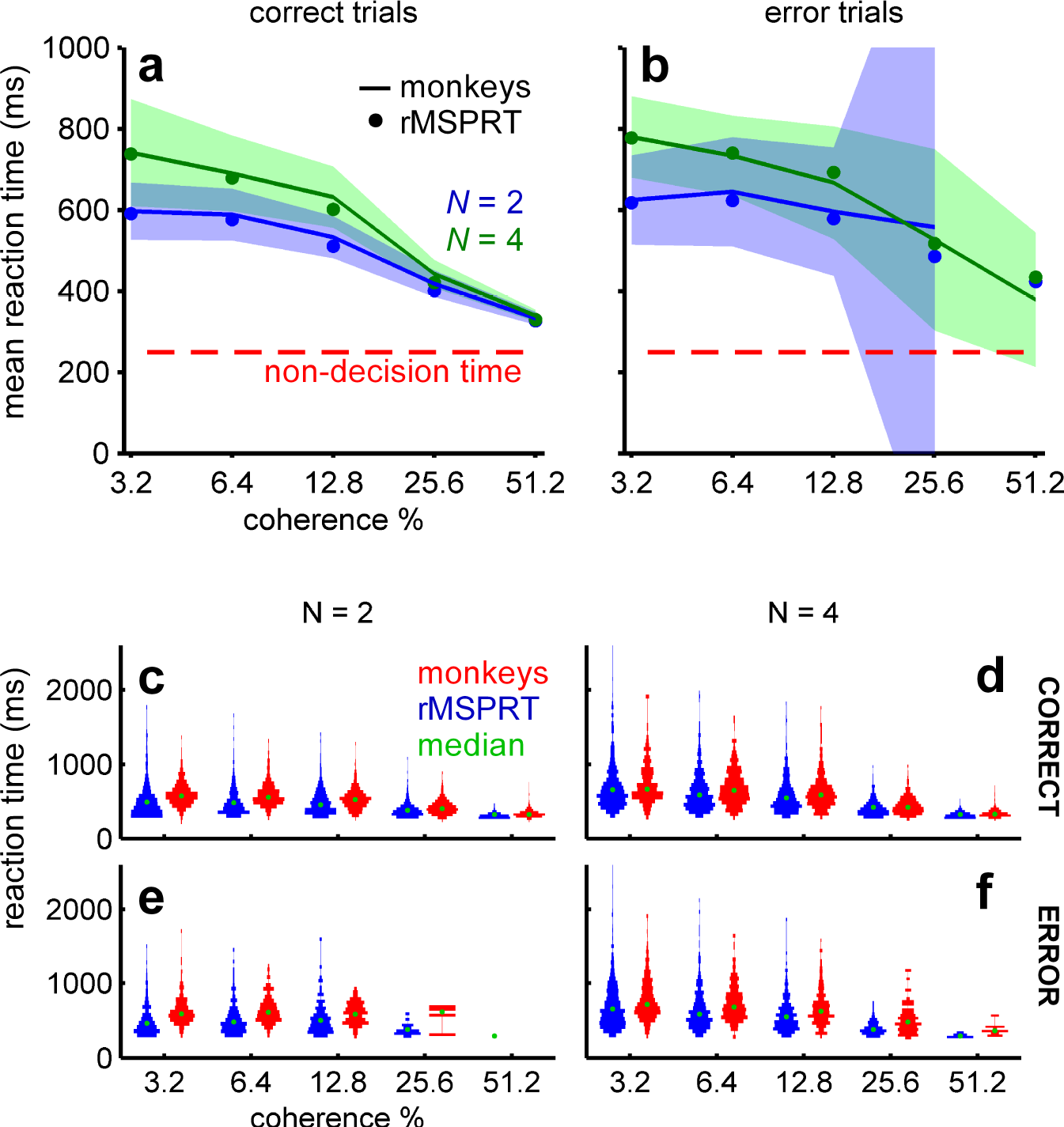
Monkey reaction times are consistent with (r)MSPRT using depleted discrimination information. (a, b) Mean reaction time of monkeys (lines) with 99 % Chebyshev confidence intervals (shading) and (r)MSPRT predictions for correct (a; Eq 14) and error trials (b; Eq 15) when using information-depleted statistics (MT parameter setΩ_*d*_). (r)MSPRT results are means of 100 simulations with 3200, 4800 total trials each for *N* = 2, 4, respectively. Confidence intervals become larger in error trials because monkeys made fewer mistakes for higher coherence levels. (c-f) ‘Violin’ plots of reaction time distributions (vertically plotted histograms reflected about the y-axis) from monkeys (red; 766-785, 1170-1217 total trials for *N* = 2, 4, respectively) and (r)MSPRT when using information-depleted statistics (blue; single Monte Carlo simulation with 800, 1200 total trials for *N* = 2,4).

### rMSPRT maps onto cortico-subcortical circuitry

The above shows that the (r)MSPRT family of exact inference algorithms can account for the dependence of choice reaction times on task difficulty, trial outcome, and the number of alternatives. But replicating behaviour alone does not tell us if the brain implements a similar computation during decisions. We thus asked whether the inner variables of the rMSPRT could account for the known dynamics of neural activity in cortex and striatum during the dot-motion task. To answer this, we must first map its components to a neural circuit. The rMSPRT is the first probabilistic model of decision able to handle recursion and arbitrary signal delays, which means that in principle it could map to a range of feedback neural circuits. Because cortex [1–5], basal ganglia [6,32] and thalamus [33] have been implicated in decision-making, we sought a mapping that could account for their collective involvement.

In the visuo-motor system, MT projects to the lateral intra-parietal area (LIP) and frontal eye fields (FEF) —two ‘sensorimotor cortex’ areas. The basal ganglia receives topographically organized afferent projections [34] from virtually the whole cortex, including LIP and FEF [35–37]. In turn, the basal ganglia provide indirect feedback to the cortex through thalamus [38,39]. This arrangement motivated the feedback embodied in rMSPRT.

Multiple parallel recurrent loops connecting cortex, basal ganglia and thalamus can be traced anatomically [38,39]. Each loop in turn can be sub-divided into topographically organised parallel loops [39,40]. Based on this, we conjecture the transient organization of these circuits into N functional loops, for decision formation, to simultaneously evaluate the possible hypotheses.

Our mapping of computations within the rMSPRT to the cortico-basal-ganglia-thalamo-cortical loop is shown in Fig 5, capturing the most prominent functional features of such circuits. For instance, it has been demonstrated that the striato-nigral and the subthalamo-nigral pathways of the basal ganglia compete during decision formation [41]. The computations predicted by rMSPRT to map on the striatum, subthalamic nucleus, and substantia nigra pars reticulata (SNr; see S3 Fig), provide a qualitative formalization of this phenomenon.

**Figure 5:**
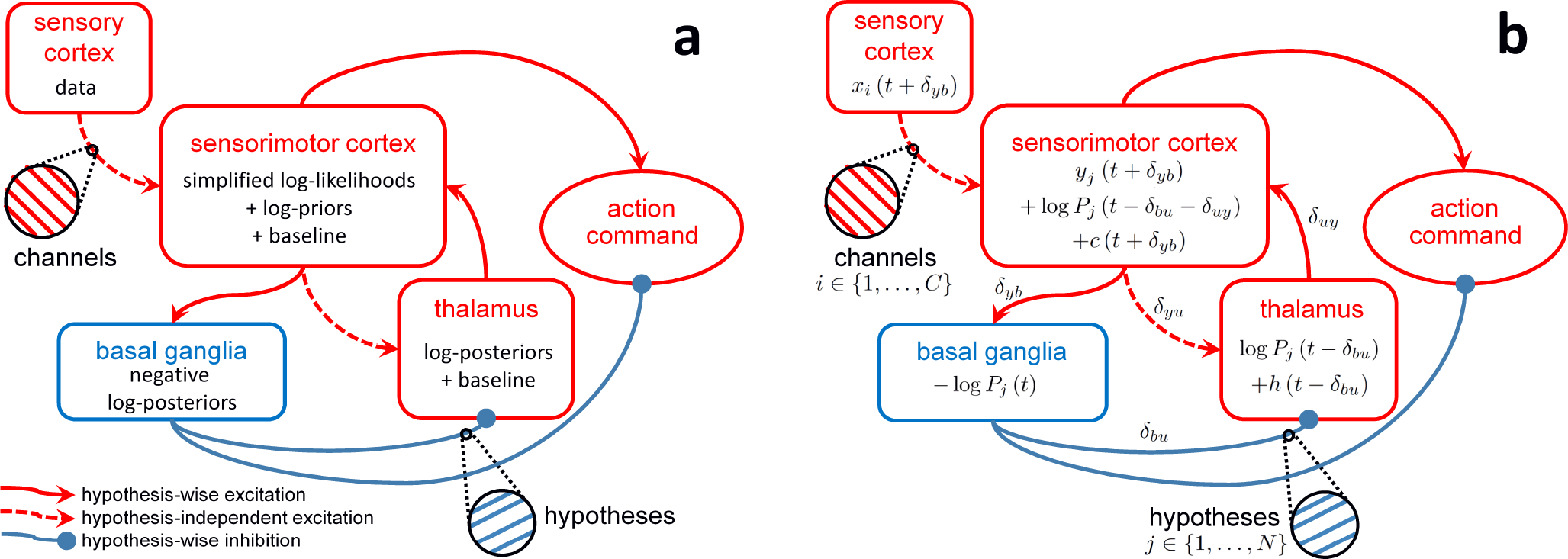
Mapping of rMSPRT computations to the cortico-basal-ganglia-thalamo-cortical loops. (a) Mapping of the negative logarithm of rMSPRT components from Fig 2. Sensory cortex (*e.g.* MT) produces fresh evidence for the decision, delivered to sensorimotor cortex in *C* parallel channels (*e.g.* MT spike trains). Sensorimotor cortex (*e.g.* LIP or FEF) computes in parallel the simplified log-likelihoods of all hypotheses given this evidence and adds log-priors —or fed-back log-posteriors after the delay Δ has elapsed. It also adds a hypothesisindependent baseline comprising a simulated constant background activity (*e.g.* from LIP before stimulus onset) and a time-increasing term from the interaction with the thalamus. The basal ganglia bring the computations of all hypotheses together into new negative log-posteriors (the output of the model basal ganglia; see S3 Fig for details) that are then tested against a threshold. Finally, the thalamus conveys the updated log-posterior from basal ganglia output to be used as a log-prior by sensorimotor cortex. Thalamus’ baseline is given by its diffuse, hypothesisindependent feedback from sensorimotor cortex. (b) Corresponding formal mapping of rMSPRT’s computational components, showing how Eq 9 decomposes. All computations are delayed with respect to the basal ganglia via the integer latencies *δ_pq_*, from *p* to *q*; where *p*, *q* ∈ {*y*, *b*, *u*}, *y* stands for the sensorimotor cortex, *b* for the basal ganglia, and u for the thalamus. Δ = *δ_yb_* + *δ_bu_* + *δ_uy_* with the requirement Δ ≥ 1.

Also, negative log-posteriors will tend to decrease for the best supported hypothesis and increase otherwise. This is consistent with the idea of basal ganglia output nuclei selectively removing inhibition from a chosen motor program while increasing inhibition of competing ones [17,32,42,43].

Lastly, our mapping of rMSPRT provides an account for the spatially diffuse cortico-thalamic projection [44], previously unaccounted for by probabilistic models of decision. It predicts that the projection conveys a constantly- increasing, hypothesis-independent baseline that does not affect the inference carried out by the cortico-basal-ganglia-thalamo-cortical loop, but may produce the offset required to facilitate the cortical re-use of inhibitory fed-back decision information from the basal ganglia (see S2 Fig). This increasing baseline may form part of the hypothesis-independent drive dubbed the “urgency signal” by [31], revealed after averaging LIP population responses across choices. All this is consistent with current views on the active modulation of information transmitted to the cortex by thalamus [45].

The mapping of rMSPRT to cortico-subcortical circuits produces key, testable predictions. First, that sensorimotor areas like LIP or FEF in the cortex evaluate the plausibility of all available alternatives in parallel, based on the evidence produced by MT, and join this to any initial bias. Second, that as these signals traverse the basal ganglia, they compete, resulting in a decision variable per alternative. Third, that the basal ganglia output nuclei use these to assess whether to make a final choice and what alternative to pick. Fourth, that decision variables are returned to sensorimotor cortex via thalamus, to become a fresh bias carrying all conclusions on the decision so far. The rMSPRT thus predicts that evidence accumulation happens uninterruptedly in the overall, large-scale loop, rather than in a single site.

### Electrophysiological comparison

With the mapping above, we can compare the dynamics of rMSPRT computations to those of recorded activity during decision-making in area LIP and striatum. We first consider the dynamics around decision initiation. During the dot motion task, the mean firing rate of LIP neurons deviates from baseline into a stereotypical dip soon after stimulus onset, possibly indicating the reset of a neural integrator [1,14]. LIP responses become choice- and coherence-modulated after the dip [1]. This also occurs when firing rates deviate from the initial baseline in striatum, where no dip is exhibited [6]. We therefore reasoned that LIP and striatal neurons engage in decision formation from the bottom of the dip or deviation from baseline (respectively) and model their mean firing rate from then on. After this, mean firing rates “ramp-up” for ∼ 40 ms in LIP, then “fork”: they continue ramping-up if dots moved towards the response (or movement) field of the neuron (inRF trials; Fig 6a, solid lines) or drop their slope if the dots were moving away from its response field (outRF trials; dashed lines) [1,3]. Striatal neurons exhibit an analogous *ramp-and-fork* pattern of response (Fig 7c,d). The magnitude of LIP firing rate is inversely proportional to the number of available alternatives (Fig 6a,b) [3,46]; a phenomenon also recorded in other visuo-motor sites, notably in the superior colliculus [47] and FEF [48–50].

**Figure 6:**
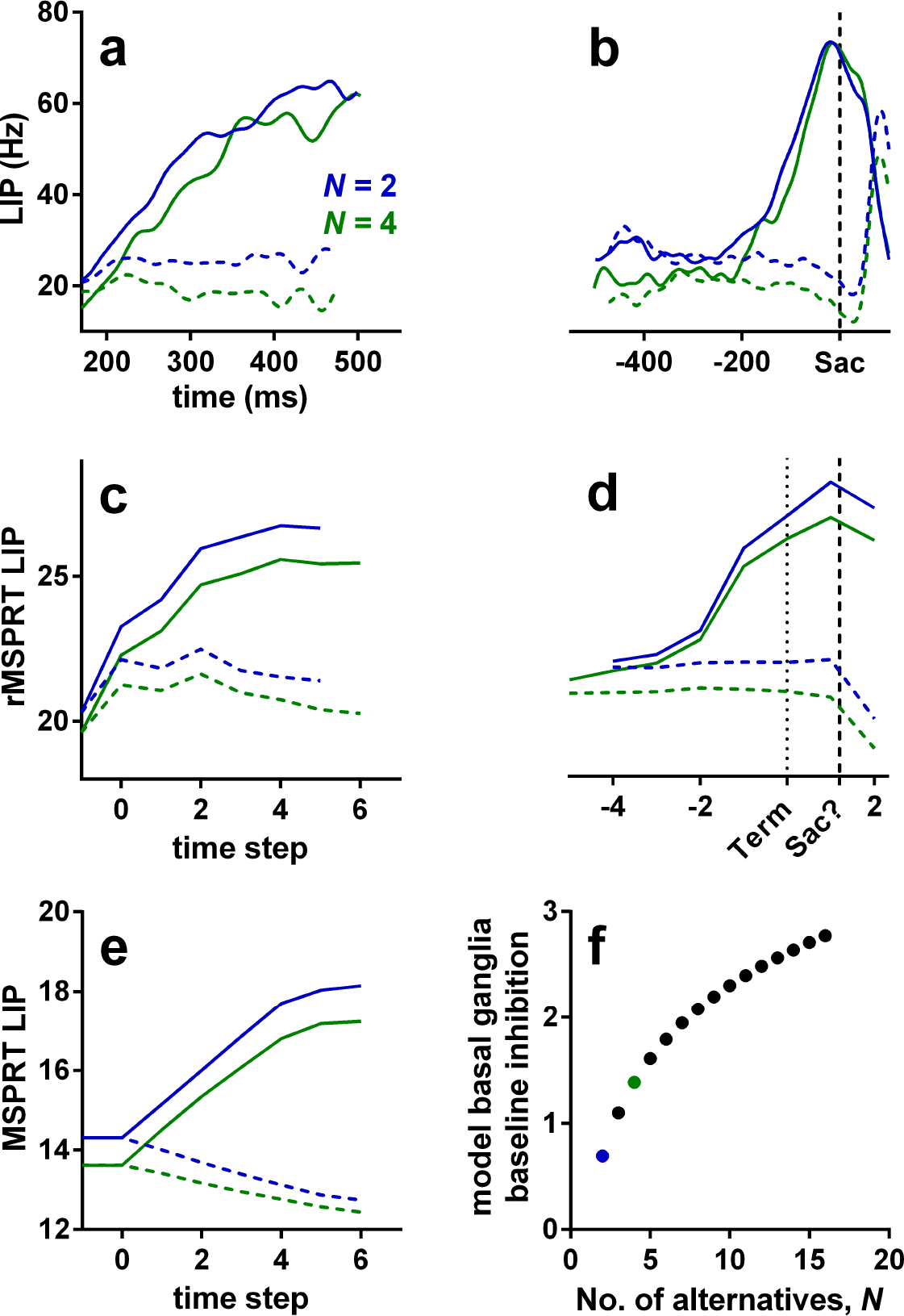
Example LIP firing rate patterns and predictions of rMSPRT and MSPRT at 25.6 % coherence. (a, b) Mean population firing rate of LIP neurons during correct trials on the reaction-time version of the dot motion task (19 neurons). By convention, inRF trials are those when recorded neurons had the motioncued target inside their response field (solid lines); outRF trials are those when that target was outside the neuron’s response field (dashed lines). (a) Aligned at stimulus onset, starting at the stereotypical dip, illustrating the “ramp- and-fork” pattern between average inRF and outRF responses. (b) Aligned at saccade onset (vertical dashed line). (c, d) Mean time course of the model sensorimotor cortex in rMSPRT aligned at decision initiation (c; *t* = 1) and termination (d; Term; dotted line), for correct trials. Initiation and termination are with respect to the time of basal ganglia output. Note the suggested saccade time “Sac?”, close to the peak of inRF computations. Simulations are a single Monte Carlo experiment with 800, 1200 total trials for *N* = 2, 4, respectively, using parameter set. For simplicity the (r)MSPRT is simulated in discrete time steps, but these have an interpretation in continous time (see Fig 9 and Methods). We include an additional step at −1 determined only by initial priors and baseline, where no inference is carried out (*y_i_* (*t* + *δ*_*yb*_) = 0 for all *i*; see Methods). Conventions as in (a). (e) Same as in (c), but for the standard, non-recursive MSPRT (Eq 10 using only the first case of Eq 8 and Eq 9). (f) Baseline output of the model basal ganglia increases as a function of the number of alternatives, thus increasing the initial inhibition of thalamus and cortex. For uniform priors, the rMSPRT predicts this function is: – log *P* (*H_i_*) = – log(1/*N*). Coloured dots indicate *N* = 2 (blue) and *N* = 4 (green).

**Figure 7:**
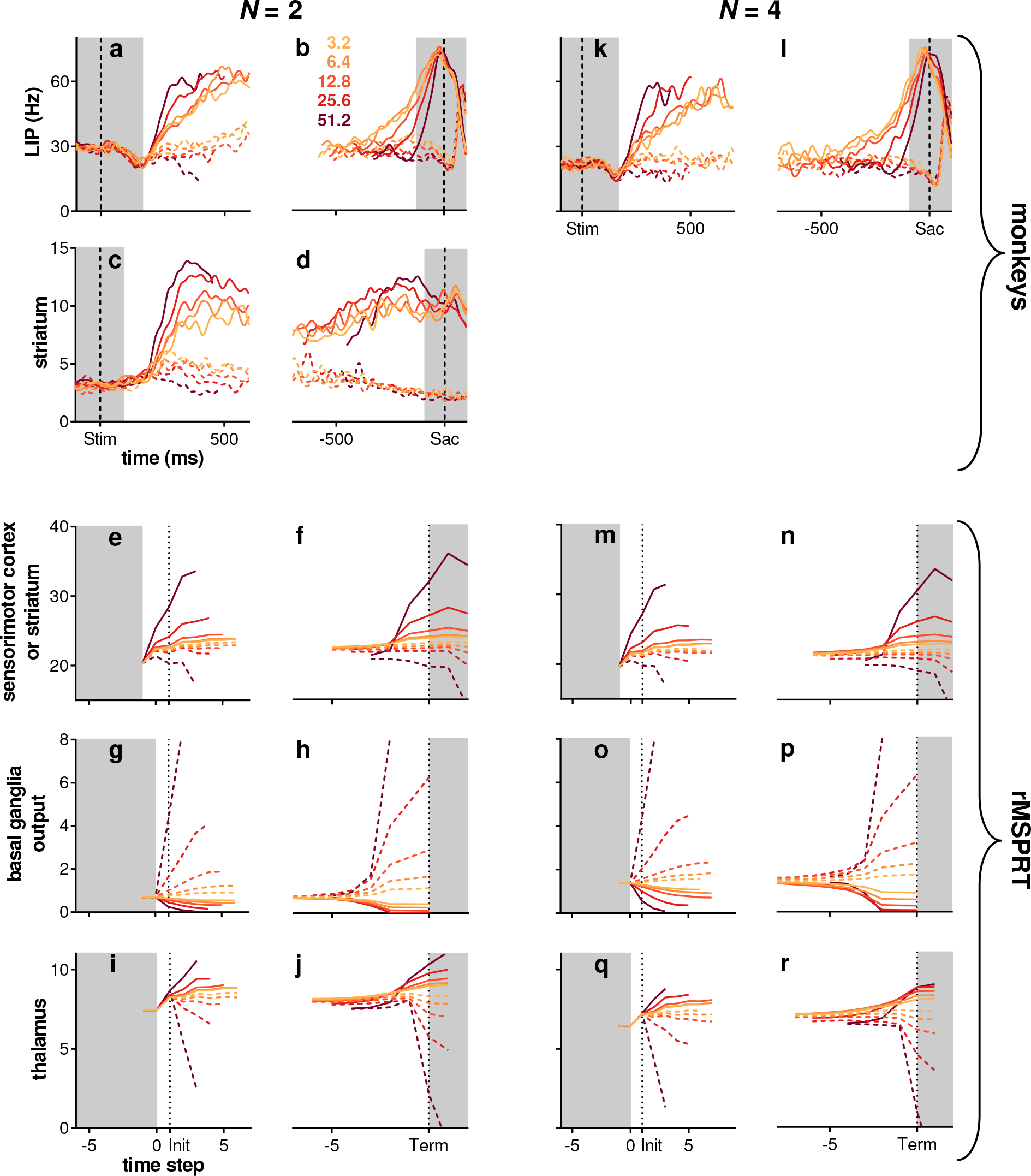
Modulation of activity by coherence throughout the cortico-basal-ganglia-thalamo-cortical loops. (a-j) *N* = 2. (k-r) *N* = 4. Top row: mean population firing rate in LIP over time during the random dot motion task (19 neurons), aligned to stimulus onset (Stim; a, k) or saccade onset (Sac; b, l) (vertical dashed lines). (c, d) mean population firing rate in striatum during the dots task (48 neurons); same conventions as in top row. (e-j, m-r) mean rMSPRT computations as mapped in Fig 5, aligned at decision initiation or termination (Init/Term; dotted lines); single Monte Carlo experiment with 800, 1200 total trials for *N* = 2, 4, respectively; simulation as in Fig 6c-e. (e, f, m, n) predicted time course of the model sensorimotor cortex (*e.g.* LIP) or striatum. (g, h, o, p) predicted simultaneous course of mean firing rate in SNr. (i, j, q, r) predicted course in thalamic relay nuclei. Solid: inRF. Dashed: outRF. Coherence % as in legend. Unshaded regions indicate approximate periods where a mechanism of decision formation should aim to reproduce the recordings.

The model LIP (sensorimotor cortex) in rMSPRT captures each of these properties: activity ramps from the start of the accumulation, forks between putative in- and out-RF responses, and scales with the number of alternatives (Fig 6c). Under this model, inRF responses in LIP occur when the likelihood function represented by neurons was best matched by the uncertain MT evidence; correspondingly, outRF responses occur when the likelihood function was not well matched by the evidence.

The rMSPRT embodies a mechanistic explanation for the ramp-and-fork pattern in the two cases of Eq 9. Initial accumulation (steps 0–2 in our simulations; feedforward inference) occurs before the feedback has arrived at the model sensorimotor cortex, resulting in a ramp. The forking (step 3; start of feedback inference) is the point at which the posteriors from the output of the model basal ganglia first arrive at sensorimotor cortex to be re-used as priors. By contrast, non-recursive MSPRT (without delayed feedback of posteriors) predicts well-separated neural signals throughout (Fig 6e). With recursion as the key difference, our framework suggests, first, that the ramp-and-fork pattern gives away the existence of an underpinning delayed inhibitory drive within a looped architecture —here from the model basal ganglia. Second, that the fork represents the time at which updated signals representing the competition between alternatives (posterior probabilities in the rMSPRT) are first made available to the sensorimotor cortex.

The rMSPRT further predicts that the scaling of activity in sensorimotor sites by the number of alternatives is due to cortico-subcortical loops becoming transiently organized as *N* parallel functional circuits, one per hypothesis. This would determine the baseline output of the basal ganglia. Until task related signals reach the model basal ganglia output, it codes the initial priors for the set of *N* hypotheses. Their output is then an increasing function of the number of alternatives (Fig 6f). This increased inhibition of thalamus in turn reduces baseline cortical activity as a function of N. The inverse proportionality of cortical activity to *N* in macaques during decisions (Fig 6a,b; [3,46,48,49]) and the direct proportionality of the firing rate to *N* in their SNr (basal ganglia output nucleus; [42]) lend support to this hypothesis.

The rMSPRT also captures key features of dynamics at decision termination. For inRF trials, the mean firing rate of LIP neurons peaks at or very close to the time of saccade onset (Fig 6b). By contrast, for outRF trials mean rates appear to fall just before saccade onset. The rMSPRT can capture both these features (Fig 6d) when we allow the algorithm to continue updating after the decision rule (Eq 10) is met. The decision rule is implemented at the output of the basal ganglia and the model sensorimotor cortex peaks just before the final posteriors have reached the cortex. The rMSPRT thus predicts that the activity in LIP lags the actual decision.

This prediction may explain an apparent paradox of LIP activity. The peri-saccadic population firing rate peak in LIP during inRF trials (Fig 6b) is commonly assumed to indicate the crossing of a threshold and thus decision termination. Visuo-motor decisions must be terminated well before saccade to allow for the delay in the execution of the motor command, conventionally assumed in the range of 80–100 ms in macaques [9,30]. It follows that LIP peaks too close to saccade onset (∼ 15 ms before) for this peak to be causal. The rMSPRT suggests that the inRF LIP peak is not indicating decision termination, but is instead a delayed read-out of termination in an upstream location.

In the rMSPRT, the striatum relays the input from sensorimotor cortex as an inhibitory drive for downstream basal ganglia nuclei. The rMSPRT has three free parameters that shape the ramp-and-fork of its inner variables, but do not alter inference. We have set their value to show that mapped variables can match the pattern in sensorimotor cortical neural dynamics (see Materials and Methods); below we show how these predictions depend on the parameter values. Nonetheless, the rMSPRT with these parameters also captures the ramp-and-fork pattern of activity in the monkey striatum (compare panels c, d to e, f in Fig 7).

LIP and striatal firing rates are also modulated by dot-motion coherence (Fig 7a-d,k,l). Following stimulus onset, the response of these neurons tends to fork more widely for higher coherence levels (Fig 7a,c,k) [1,3,6]. The increase in activity before a saccade during inRF trials is steeper for higher coherence levels, reflecting the shorter average reaction times (Fig 7b,d,l) [1,3,6]. The sensorimotor cortex or striatum in the rMSPRT shows coherence modulation of both the forking pattern (Fig 7e,m) and slope of activity increase (Fig 7f,n). rMSPRT also predicts that the apparent convergence of peri-saccadic LIP activity to a common level during inRF trials (Fig 7b,l) is not required for inference and so may arise due to additional neural constraints. We take up this point in the Discussion.

### Electrophysiological predictions

Our proposed mapping of the rMSPRT’s components (Fig 5) makes testable qualitative predictions for the mean responses in basal ganglia and thalamus during the dot motion task. For the basal ganglia output, likely from the oculomotor regions of the SNr, rMSPRT (like MSPRT) predicts a drop in the activity of output neurons during inRF trials and an increase in outRF ones. It also predicts that these changes are more pronounced for higher coherence levels (Fig 7g,h,o,p). These predictions are consistent with recordings from macaque SNr neurons showing that they suppress their inhibitory activity during visually- or memory-guided saccade tasks, in putative support of saccades towards a preferred region of the visual field [42,51,52], and enhance it otherwise [52].

In detection tasks like visually- or memory-guided ones, the decision cues are extremely obvious. Hence, the accompanying recorded neural-activity transients may be argued to encode very short evidence-accumulations. After all, the accumulation of a single observation (*e.g.* an ISI) is the simplest, albeit degenerate case of evidence accumulation.

For visuo-motor thalamus, rMSPRT predicts that the time course of the mean firing rate will exhibit a ramp-and-fork pattern similar to that in LIP (Fig 7i,j,q,r). The separation of in- and out-RF activity is consistent with the results of [33] who found that, during a memory-guided saccade task, neurons in the macaque medio-dorsal nucleus of the thalamus (interconnected with LIP and FEF), responded more vigorously when the saccade target was flashed within their response field than when it was flashed in the opposite location.

### Predictions for neural activity features not crucial for inference

Understanding how a neural system implements an algorithm is complicated by the need to identify which features are core to executing the algorithm, and which are imposed by the constraints of implementing computations using neural elements —for example, that neurons cannot have negative firing rates, so cannot straightforwardly represent negative numbers. The three free parameters in the rMSPRT allow us to propose which functional and anatomical properties of the cortico-basal-ganglia-thalamo-cortical loop are workarounds within these constraints, but do not affect inference.

One free parameter enforces the baseline activity that LIP neurons maintain before and during the initial stimulus presentation (Fig 7a,k). Varying this parameter, *l*, scales the overall activity of LIP, but does not change the inference performed (Fig 8a). Consequently, this suggests that the baseline activity of LIP depends on *N* but does not otherwise affect the inference algorithm implemented by the brain.

**Figure 8:**
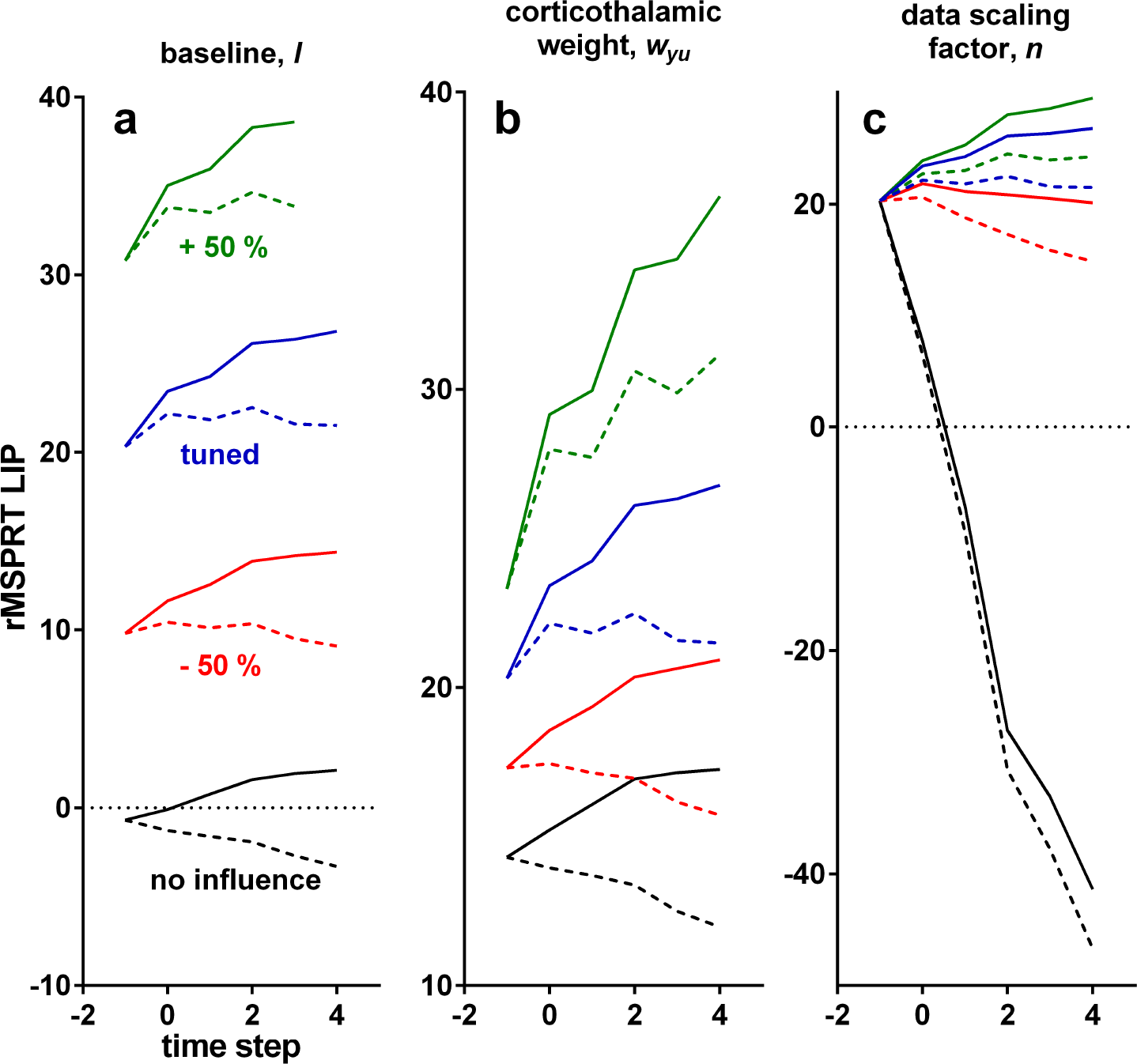
Effect of variations of free parameters on the time course of the model LIP in rMSPRT. Each solid and dashed set of lines is the mean of correct trials in a single Monte Carlo experiment, with 800 total trials, 25.6 % coherence and *N* = 2; simulation as in Fig 6c-e. Computations aligned at decision initiation. Solid: inRF. Dashed: outRF. Blue: with parameters as tuned for this study. Green: increasing parameter value by 50 %, keeping other parameters as tuned. Red: decreasing it by 50 %, keeping others as tuned. Black: removing the effect of the tested parameter (*l* = 0, *w_yu_* = 0, *n* = 1), keeping others as tuned. (a) Varying the baseline, *l*. (b) Varying the cortico-thalamic weight, *w_yu_*. (c) Varying the data scaling factor, *n*.

The second free parameter, *w_yt_*, sets the strength of the spatially diffuse projection from cortex to thalamus. Varying this weight changes the forking between inRF and outRF computations but does not affect inference (Fig 8b). The third free parameter, *n*, sets the overall, hypothesis-independent temporal scale at which sampled input ISIs are processed; changing *n* varies the slope of sensorimotor computations, even allowing all-decreasing mean firing rates (Fig 8c). By definition, the log-likelihood of a sequence tends to be negative and decreases monotonically as the sequence lengthens. Introducing n is required to get positive simplified log-likelihoods, capable of matching the neural activity dynamics, without affecting inference. Hence, *n* may capture a workaround of the decision-making circuitry to represent these whilst avoiding signal ‘underflow’, by means of scaling the input data.

Traditionally, evidence accumulation is exclusively associated with increasing firing rates during decision, and previous studies have questioned whether the often-observed decision-correlated yet non-increasing firing rates (*e.g.* in outRF conditions in Fig 7a,c,k and [1–3,5,53,54]) are consistent with accumulation [22,23]. The diversity of patterns predicted by rMSPRT in sensorimotor cortex (Fig 8) solves this by demonstrating that both increasing and non-increasing activity patterns can house evidence accumulation.

**Figure 9:**
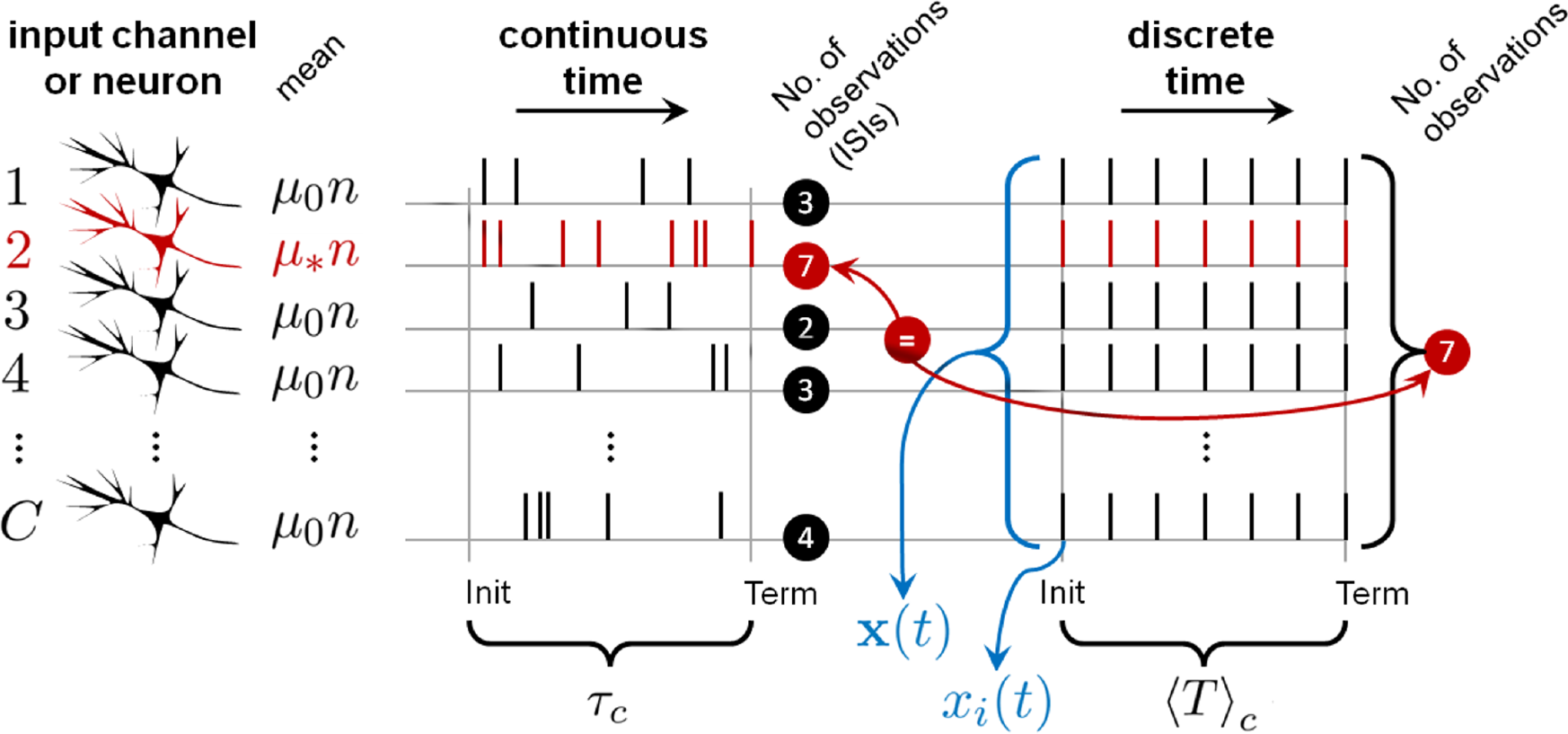
The mean number of ISIs to decision in continuous-time spike-trains is equivalent to the mean number observations to decision in discrete time. C sensory neurons (input channels; left) produce sequences of ISIs in continuous time with mean μ*n (red; best tuned to the stimulus) or *μ_0_n* (black; otherwise). The average decision time —between decision initiation (Init) and termination (Term)— is **α**_*C*_ in correct trials, as in this diagram. In discrete time it takes an average of 〈T〉_c_ vector observations, **x**(*t*) (composed of scalar observations *x_i_*(*t*), each time step *t*; blue), to make decisions. [20] showed that in the minimum input case (when *C* = *N*), the mean number of ISIs in the most active channel (red) used by a general, continuous-time, spike-based instance of the MSPRT, approximately equal the mean number of observations, 〈T〉_c_, required by the simpler, discrete-time MSPRT (here 7 in both cases), which carries to our identically-performing rMSPRT; this is true under equal input channel statistics (μ_*_, μ_0_, σ_*_, σ_0_), data distributions (*e.g.* all lognormal), number of alternatives, *N*, and error rate, e. This all implies that, if we add 0.5 —the expected number of ISIs from decision initiation to a first spike— to 〈T〉_c_, and multiply this by the minimum mean ISI, *μ*_*_*n* (of fastest firing channel), this approximately equals τ_*c*_, hence Eq 14; conversely, in error trials we use *μ*_0_ and 〈T〉_e_, to get τ_e_ (Eq 15).

## Discussion

We tested the hypothesis that the brain approximates exact inference for decision making. We did so by showing that a novel recursive form of the MSPRT, the rMSPRT, uniquely accounts for both monkey choice behaviour and the corresponding neural dynamics in cortex and striatum, while its architecture matches that of the cortico-subcortical decision circuits.

### Why implement a recursive procedure in the brain?

The recursive computation implied by the looped cortico-basal-ganglia-thalamo-cortical architecture has several advantages over local or feedforward computations. First, recursion makes trial-to-trial adaptation of decisions possible. Priors determined by previous stimulation (fed-back posteriors), can bias upcoming similar decisions towards the expected best choice, even before any new evidence is collected. This can shorten reaction times in future familiar settings without compromising accuracy. Second, recursion provides a robust memory. A posterior fed-back as a prior is a sufficient statistic of all past evidence observations. That is, it has taken ‘on-board’ all sensory information since the decision onset. In rMSPRT, the sensorimotor cortex only need keep track of observations in a moving time window of maximum width Δ —the delay around the cortico-subcortical loop- rather than keeping track of the entire sequence of observations. For a physical substrate subject to dynamics and leakage, like a neuron in LIP or FEF, this has obvious advantages: it would reduce the demand for keeping a perfect record (*e.g.* likelihood) of all evidence, from the usual hundreds of milliseconds in decision times to the ∼ 30 ms of latency around the cortico-basal-ganglia-thalamo-cortical loop (adding up estimates from [55–57]).

### Lost information and perfect integration

The rMSPRT decides faster than monkeys in the same conditions because monkeys do not make full use of the discrimination information available in their MT (Fig 3b). However, this performance gap arises partially because rMSPRT is a generative model of the task. Thus, this assumes that knowledge of coherence is available by decision initiation, which in turn determines appropriate likelihoods for the task at hand. Any deviation from this generative model will tend to degrade performance, whether it comes from one or more of: the coherence to likelihood mapping [58], the inherent leakiness of neurons, or correlations between spikes or between neurons (see [20]). In this respect, we must consider, first, that the activity dip ∼ 170 ms after stimulus onset is assumed to indicate decision engagement at the LIP level. By then, MT neurons have been reliably modulated by motion coherence for about 120 ms (starting ∼ 50 ms after stimulus onset; see S4 Fig for details), giving a sizeable window to adjust LIP ‘likelihood functions’ to match the decision at hand. Whether this window is large enough or if trial-by-trial ‘likelihood adjustment’ occurs at all remain as interesting questions for future experimental explorations. Second, that LIP neurons change their coding during learning of the dot motion task and MT neurons do not [59], implying that learning the task requires mapping of MT to LIP populations by synaptic plasticity [60]. Consequently, even if the MT representation is perfect, the learnt mapping only need satisfice the task requirements, not optimally perform.

Excellent matches to monkeys’ performance in both correct and error trials, and hence their speed-accuracy trade-offs, were obtained solely by accounting for lost information in the evidence streams. No noise was added within the rMSPRT itself. Prior experimental work reported perfect, noiseless evidence integration by both rat and human subjects performing an auditory task, attributing all effects of noise on task performance to the variability in the sensory input [61]. Our results extend this observation to primate performance on the dot motion task, and further support the idea that the neural decision-making mechanism can perform perfect integration of uncertain evidence.

### Neural response patterns during decision formation

Neurons in LIP, FEF [4], and striatum exhibit a ramp-and-fork pattern during the dot motion task. Analogous choice-modulated patterns have been recorded in the medial premotor cortex of the macaque during a vibro-tactile discrimination task [53] and in the posterior parietal cortex and frontal orienting fields of the rat during an auditory discrimination task [5]. The rMSPRT indicates that such slow dynamics emerge from decision circuits with a delayed, inhibitory drive within a looped architecture. This suggests that decision formation in mammals may use a common recursive computation.

A random dot stimulus pulse delivered earlier in a trial has a bigger impact on LIP firing rate than a later one [2]. This highlights the importance of capturing the initial, early-evidence ramping-up before the forking. However, multiple models omit it, focusing only on the forking (*e.g.* [9,10, 13]). Other, heuristic models account for LIP activity from the onset of the choice targets, through dots stimulation and up until saccade onset (*e.g.* [12,14–16]). Nevertheless, their predicted firing rates rely on two fitted heuristic signals that shape both the post-stimulus dip and the ramp-and-fork pattern. In contrast, the ramp-and-fork dynamics emerge naturally from the delayed inhibitory feedback in rMSPRT during decision formation.

rMSPRT qualitatively replicates the ramp-and-fork pattern for individual coherence levels and given number of alternatives, *N* (Fig 6). However, the peak of the accumulated evidence in the model sensorimotor cortex of rMSPRT does not converge to a common value around decision termination during inRF trials. Consequently, it predicts that the apparent convergence of LIP activity to a common value (Fig 6b, Fig 7b,l) is not part of the inference procedure, but reflects other constraints on neural activity.

One such constraint is that these brain regions engage in multiple other computations, some of which are likely orthogonal to solving the random dot motion task. The neural activity recorded during decision tasks may then be a transformation of inference computations, by mixing them with all other simultaneous computations. Consistent with this, the successful fitting of previous computational models to neural data [12, 14–16] has been critically dependent on the addition of heuristic signals for unknown constraints. While beyond the scope of this study, which examined whether a normative mechanism could explain behaviour and electrophysiology during decisions, adding similar heuristic signals to the rMSPRT would likely allow a quantitative reproduction of the peri-saccadic convergence of LIP activity.

### Emergent predictions

Inputs to the rMSPRT were determined solely from MT responses during the dot-motion task, and it has only three free parameters, none of which affect inference. It is thus surprising that it renders emergent predictions that are consistent with experimental data. First, our information-depletion procedure used exclusively statistics from correct trials. Yet, after depletion, rMSPRT matches monkey behaviour in correct *and* error trials (Fig 4), suggesting a mechanistic connection between them in the monkey that is naturally captured by rMSPRT. Second, the values of the three free parameters were chosen solely so that the model LIP activity resembled the ramp-and-fork pattern observed in our LIP data-set (Fig 6a,c). As demonstrated in Fig 8, the ramp-and-fork pattern is a particular case of two-stage patterns that are an intrinsic property of the rMSPRT, guaranteed by the feedback of the posterior after the delay Δ has elapsed (Eq 5). Nonetheless, the algorithm also qualitatively matches LIP dynamics when aligned at decision termination (6b,d). Third, the predictions of the time course of the firing rate in SNr and thalamic nuclei naturally emerge from the functional mapping of the algorithm onto the cortico-basal-ganglia-thalamo-cortical circuitry. These are already congruent with existing electrophysiological data; however, their full verification awaits recordings from these sites during the dot motion task. These and other emergent predictions are an encouraging indicator of the explanatory power of a systematic framework for understanding decision formation, embodied by the rMSPRT.

### Relation of the rMSPRT to prior decision models

The rMSPRT contains all previous instances of the MSPRT [17,18,25,26,62] as special cases. It generalizes them by allowing the re-use of posteriors at any given time in the past as priors for present inference, via recursion. The (r)MSPRT also contains the sequential probability ratio test when *N* = 2, and its continuous-time equivalent, the popular drift-diffusion model (*e.g.* [4,6,9,61,63–66]). While a valuable basic model of decision-making, the drift-diffusion model is restricted to *N* = 2 alternatives and does not address neural mechanisms. First, it assumes that evidence for decisions comes as a continuous Gaussian process whose presence in the brain is unproven. Since the decision times predicted by the model critically hinge on this process and its statistics (typically disconnected from the statistics of sensory neural activity), this limitation also obscures the interpretation of the drift-diffusion model’s behavioural predictions. Second, its single decision variable must restrict itself to the half-plane closest to the choice threshold associated to one of its two hypotheses if such hypothesis is to be chosen; hence, the drift-diffusion model can account for forking dynamics, but not for the preceding ramping observed in experimental data. In contrast, the rMSPRT natively captures decisions among any number of alternatives (N ≥ 2), can explain ramp-and-fork dynamics, and does so using neural evidence for decisions in its natural format: spike-trains with statistics extracted from MT recordings.

Biophysical models that directly address neural implementations of decision making are predominantly based on winner-take-all competition between neurons representing different hypotheses [8,11–14,16,67,68]. These provide valuable insights into potential mechanisms by which neural circuits can represent and compute decisions, but do not typically make contact with formal inference procedures (see [69]). The studies of [13,68] are possible exceptions, since they make the analogy between the predictions of their neural-network model and those of exact, Bayes-based inference. Conversely, the rMSPRT shows how a normative decision-making algorithm can account for cortical and subcortical activity. As such, the rMSPRT provides target, exact-inference computations for future biophysical models.

### Anatomical mapping, assumptions, and future directions

Mapping any formal algorithm to a neural substrate implies proposing assumed computational contributions for the components of the substrate. In mapping the rMSPRT we made two broad classes of assumptions. First, as explained above, that individual substrates implement multiple functions either simultaneously or under different stimulation scenarios (*e.g.* experimental paradigms). In particular, we assume that during decision-formation the striatum is only required to perform a light-touch, relay-like transformation of its excitatory cortical inputs into inhibitory outputs. This assumption is shared by multiple models of the basal ganglia (*e.g.* [70,71]). The similarity between ramp-and-fork patterns of response across neurons in the LIP [3], FEF [4], and striatum [6] during the dot-motion task, is consistent with this (Fig 7a-d). That said, computational models have shown how the striatum’s intricate microcircuit [72] can give rise to several types of complex responses to simple cortical input, often taking the form of spontaneously appearing neural ensembles [73–75]. Thus a promising avenue for future research is determining if, and how, the dynamics of the striatal micro-circuit can act as a relay-like function during decision formation.

Our second class of assumptions is that the omitted connections into and within the basal ganglia may not contribute to the computations essential to inference with cortical inputs. Of note, we have omitted in our mapping the projections from thalamus to striatum [76] or to subthalamic nucleus [77], as well as the intrinsic connections from subthalamic nucleus or from globus pallidus pars externa (globus pallidus in non-primates) to striatum (*e.g.* see [77,78]). Such omitted connections might offer a more robust implementation of inference computations, or may contribute to overcoming the limitations of implementing an algorithm with neurons.

Demonstrating the compatibility of anatomical pathways with the mapping of the (r)MSPRT is the subject of ongoing research. Success has been achieved in the expansion of the basal-ganglia mapping of the MSPRT to include the pathway from striatum to globus pallidus pars externa and that from the latter to SNr, where the same inference could be done without those pathways [17]. It has also been recently shown that the pallido-striatal connection is compatible with the MSPRT mapping onto the basal ganglia [21], possibly giving a more robust neural implementation (both results carry to the rMSPRT). In the same bracket is our inclusion of the cortico-thalamic projection here (Fig 5). Since this projection is assumed to be hypothesis-independent (Eq 12), it does not affect the inference done by the rMSPRT. Similar exercises may be able to account for projections from thalamus to striatum or to subthalamic nucleus, and from the latter to striatum, though these are beyond the scope of this study. The rMSPRT provides a starting point to explore all such extended mapping alternatives.

## Conclusion

We sought to characterize the neural mechanism that underlies decisions using a normative algorithm —the rMSPRT— as a framework. We find it remarkable that, starting from data-constrained spike-trains, our monolithic statistical test can simultaneously account for much of the anatomy, behaviour, and electrophysiology of decisionmaking. While it is not plausible that the brain implements exactly a specific algorithm, our results suggest that the essential composition of its underlying decision mechanism includes the following. First, that the mechanism is probabilistic in nature —the brain utilizes the uncertainty in neural signals, rather than suffering from it. Second, that the mechanism works entirely ‘on-line’, continuously updating representations of hypotheses that can be queried at any time to make a decision. Third, that this processing is distributed, recursive, and parallel, producing a decision variable for each available hypothesis. And fourth, that this recursion allows the mechanism to adapt to the observed statistics of the environment in an unsupervised manner, as it can re-use updated probabilities about hypotheses as priors for upcoming decisions. With the currently available range of experimental studies giving us local snapshots of cortical and subcortical activity during decision-making tasks, the rMSPRT shows us how, where, and when these snapshots fit into a complete inference procedure.

## Materials and Methods

### Experimental paradigms

Behavioural and neural data was collected in three previous studies [3,6,24], during two versions of the random dot motion task (Fig 1a-c). Detailed experimental protocols can be found in each report. Below we briefly summarize them.

#### Fixed duration

Three rhesus macaques (*Macaca mulatta*) were trained to initially fixate their gaze on a visual fixation point (cross in Fig 1a). A random dot kinematogram appeared covering the response field of the MT neuron being recorded (grey patch); task difficulty was controlled per trial by the proportion of dots (coherence %) that moved in one of two directions: that to which the MT neuron was tuned to —its preferred motion direction— or its opposite —null motion direction. After 2 s the fixation point and kinematogram vanished and two targets appeared in the possible motion directions. Monkeys received a liquid reward if they then saccaded to the target towards which the dots in the stimulus were predominantly moving [24].

#### Reaction time

Two macaques per study learned to fixate their gaze on a central fixation point (Fig 1b,c). Two (Fig 1b) or four (Fig 1c; only in the protocol of [3]) eccentric targets appeared, signalling the number of alternatives in the trial, *N*. One such target fell within the response (movement) field of the recorded neuron (grey patch). This is the region of the visual field towards which the neuron would best support a saccade. Later a random dot kinematogram appeared where a controlled proportion of dots moved towards one of the targets. The monkeys received a liquid reward for saccading to the indicated target when ready [3,6].

### Data analysis

For comparability across databases, we only analysed data from trials with coherence levels of 3.2, 6.4, 12.8, 25.6, and 51.2 %, unless otherwise stated. We used data from all neurons recorded in such trials. Our datasets contained between 189 and 213 visual-motion-sensitive MT neurons (see Table 1; single-cell recordings from [24,79]), as well as 19 LIP neurons (data from [3]) and 48 striatal ones (from [6]) whose activity was previously determined to be choice- and coherence-modulated. The behavioural data analysed was that associated to LIP recordings. For MT, we analysed the neural activity between the onset and the vanishing of the stimulus. For LIP and striatum we focused on the period between 100 ms before stimulus onset and 100 ms after saccade onset.

**Table 1:** Population ISI statistics (ms) in MT per coherence (first column).

| Coherence % | No. neurons | $\Omega, \Omega_d$ | | $\Omega$ | | $\Omega_d, N = 2$ | | $\Omega_d, N = 4$ | |
| --- | --- | --- | --- | --- | --- | --- | --- | --- | --- |
| | | $\mu_*n$ | $\sigma_*n$ | $\mu_0n$ | $\sigma_0n$ | $\mu_0n$ | $\sigma_0n$ | $\mu_0n$ | $\sigma_0n$ |
| 3.2 | 206 | 54.1 | 33.1 | 59.4 | 34.5 | 59.0 | 34.4 | 58.5 | 34.3 |
| 6.4 | 211 | 52.0 | 32.2 | 62.9 | 35.3 | 60.6 | 34.7 | 59.8 | 34.4 |
| 12.5 | 213 | 46.1 | 30.5 | 65.5 | 36.1 | 60.3 | 34.6 | 58.3 | 34.0 |
| 25.5 | 208 | 37.7 | 28.0 | 70.2 | 37.2 | 62.0 | 34.9 | 59.9 | 34.3 |
| 51.2 | 189 | 29.9 | 26.0 | 83.5 | 40.6 | 75.5 | 38.5 | 71.8 | 37.4 |

To estimate moving statistics of neural activity we first computed the spike count over a 20 ms window sliding every 1 ms, per trial. The moving mean firing rate per neuron per condition was then the mean spike count over the valid bins of all trials divided by the width of this window; the standard deviation was estimated analogously. LIP and striatal recordings were either aligned at the onset of the stimulus or of the saccade; after or before these (respectively), data was only valid for a period equal to the reaction time per trial. The population moving mean firing rate is the mean of single-neuron moving means over valid bins; analogously, the population moving variance of the firing rate is the mean of single neuron moving variances. For clarity, population statistics were then smoothed by convolving them with a Gaussian kernel with a 10 ms standard deviation. The resulting smoothed population moving statistics for MT are in Fig. 1d,e. LIP and striatal mean firing rates are plotted only up to the median reaction time plus 80 ms, per condition.

Analogous procedures were used to compute the moving mean of the computations within simulated algorithms, per time step, rather than over a moving window. These are shown up to the median of termination observations plus 3 time steps.

### Definition of the recursive multi-hypothesis sequential probability ratio test (rM-SPRT)

Let x(*t*) = (*x_i_* (*t*),…,*x_c_* (*t*)) be a vector random variable composed of scalar observations, *x_j_* (*t*), made in *C* channels at time *t* ∈ {1, 2,…} (right-hand side of Fig 9). Let also x(*r*:*t*) = (x(*r*)/*n*,…, x(*t*)/*n*) be the sequence of vectors x(*t*)/*n*, i.i.d. across time, from r to t (r < t). Here *n* ∈ {ℝ > 0} is a constant data scaling factor. If *n* > 1, it scales down incoming data, *X_j_* (*t*); this will prove useful ahead when tuning the algorithm to reveal that the dynamics in rMSPRT computations match those of sensorimotor cortex. Note that scaling is only effective from the likelihood on and does not affect the time interpretation of the data. Crucially, since n is hypothesis-independent, it does not affect inference.

There are *N* ∈{2, 3,…} alternatives or hypotheses about the uncertain *evidence*, x(1: *t*) —say possible courses of action or perceptual interpretations of sensory data. The task of a decision maker is to determine which hypothesis *H_i_* (*i* ∈ {1,…, *N*}) is best supported by this evidence as soon as possible, for a given level of accuracy. To do this, it requires to estimate the posterior probability of each hypothesis given the data, *P* (*H*_*i*_|x(1: *t*)), as formalized by Bayes’ rule. The mechanism we seek must be recursive to match the nature of the brain circuitry. Formally, *P* (*H*_*i*_|x(1: *t*)) will be initially computed upon starting priors *P* (*H*_*i*_) and likelihoods *P*(x(1: *t*)|*H*_i_); however, after some time Δ ∈ {1, 2,…}, it will re-use past posteriors, *P* (*H*_*i*_|x(1: *t* – Δ)), Δ time steps ago, as priors, along with the likelihood function *P* (x(*t* – Δ + 1: *t*)|*H*_*i*_) of the segment of x(1: *t*) not yet accounted by *P* (*H*_i_|x(1: *t* – Δ)). A mathematical induction proof of this form of Bayes’ rule follows.

If say Δ = 2, in the first time step, *t* = 1:

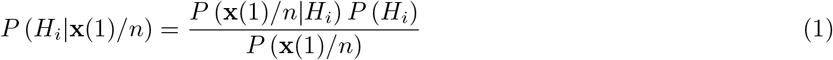

By *t* = 2:

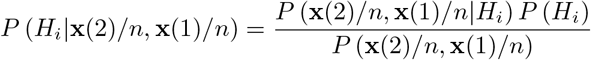

Note that we are still using the initial fixed priors *P* (*H*_*i*_). Now, for *t* = 3:

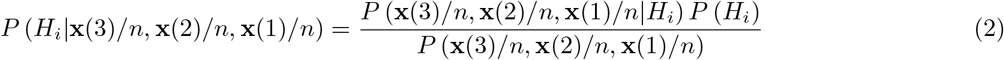

According to the product rule, we can segment the probability of the sequence x(1: *t*) as:

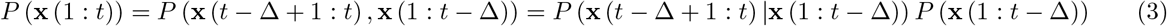

And, since x(*t*) are i.i.d., the likelihood of the two segments is:

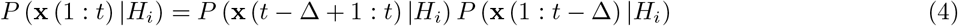

If we substitute the likelihood in Eq 2 by Eq 4, its normalization constant by Eq 3 and re-group, we get:

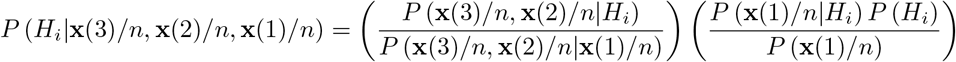

It is evident that the rightmost factor is *P*(*H_i_*\**x**(1)/*n*) as in Eq 1. Hence, in this example, by *t* = 3 we start using past posteriors as priors for present inference as:

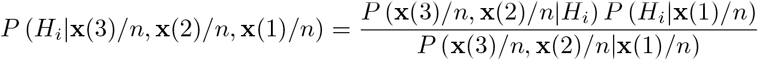

So, in general:

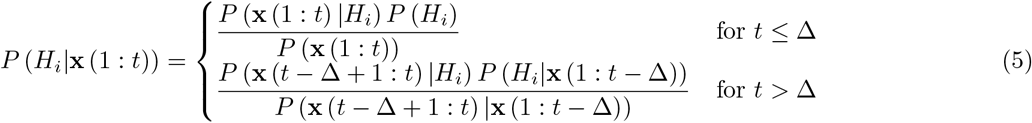

where the normalization constants are

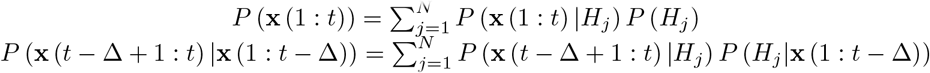

Eq 5 is a general recursive form of the Bayes’ rule, designed to accumulate evidence for inference in a recurrent, uninterrupted fashion. By *t* > Δ, it uses posteriors Δ ≥ 1 time steps in the past as current priors, thereby generalizing a previous common recursive form of the Bayes’ rule that is limited to Δ = 1 (that in *e.g.* [18, 26, 68, 80,81]). Priors updated in this manner are a sufficient statistic of all the evidence observed up to *t* – Δ. By this ability, and in the general machine-learning sense, any decision algorithm harnessing Eq 5 adapts or learns. Since no labelled examples or teaching signals are required for such learning, the rMSPRT is thence said to be engaged in ongoing *unsupervised learning.*
Ahead we use three key results from [20] as part of our methods, with no overlap between their results and the results of the present study. First, a lognormal-based form of the likelihood function whose component operations they showed are neurally plausible and most consistent with the statistics of MT responses during the random dots task. Second, a crucial link between the statistics of ISIs in the spike-trains used as evidence for decision (*e.g.* those of MT during the dots task), and continuously-distributed MSPRT decision times. As discussed below, this link enabled us to use simpler, discrete-time algorithms and still interpret their behavioural predictions in continuous time. And third, the fundamental dependence of MSPRT decision times on: (a) the discrimination information available in the evidence and (b) a constant, fixed for given error rate and *N*. Since rMSPRT performs identically to MSPRT, all this carries to it.

It is apparent that the critical computations in Eq 5 are the likelihood functions. The forms that we consider ahead build upon the simplest shown by [20], where the number of evidence streams equals the number of hypotheses (*C* = *N*); for instance, a minimum of *C* = 2 differently-tuned neurons are assumed to provide evidence for a *N* = 2 choice decision. As discussed by them, more complex (*C* > *N*), biologically-plausible likelihood functions can be formulated if necessary; the *C* < *N* case would make no sense as it would imply the testing of redundant hypotheses. Although not essential, to simplify the notation when C = *N*, from now on data in the channel conveying the most salient evidence for hypothesis *H_i_* will bear its same index *i*, as *x_i_* (*j*). When *t* ≤ Δ we have:

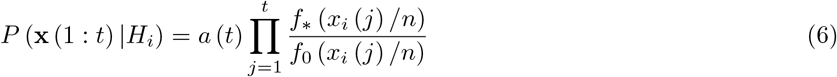

this is, the likelihood that *x*_i_ (*j*)/*n* was drawn from a distribution, *f*_*_, rather than from *f*_0_, that is assumed to have originated *x*_*k*_ (*j*)/*n* (*k* ≠ *i*) for the rest of the channels. In Eq 6, 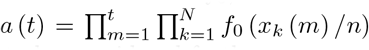 is a hypothesis-independent factor that does not affect Eq 5 and thus needs not to be considered further.

When *t* > Δ only the observations in the time window [*t* – Δ + 1, *t*] are used for the likelihood function because data before this window is already considered within the fed-back posterior, *P* (*H*_*i*_|x(1: *t* – Δ)). Then, the likelihood function is:

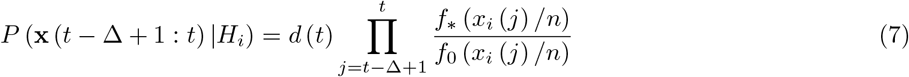

where again 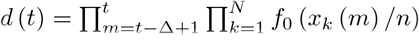 needs not to be considered further.

Now, for our likelihood functions to work upon a statistical structure like that produced by neurons in MT we need to be more specific. Inter-spike intervals (ISI) in MT during the random dot motion task are best described as lognormally distributed [20] and we assume that decisions are made upon the information conveyed by them. Thus, from now on we assume that *f*_*_ and *f*_0_ are lognormal and that they are specified by means μ_*_ and μ_0_, and standard deviations σ_*_ and σ_0_, respectively. We can then put together the logarithm of Eq 6 and Eq 7 as the log-likelihood function, *y_i_* (*t*), substituting the lognormal-based form of it reported by [20]:

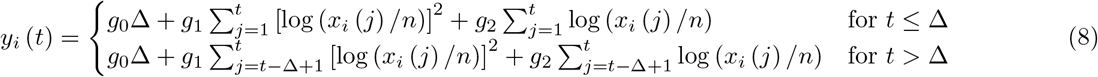

with

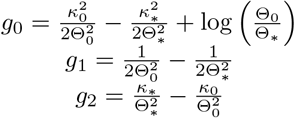

where 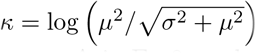 and θ^2^ = (σ^2^/μ^2^ + l) with appropriate subindices *, o.

The terms *g*_0_Δ in Eq 8 are hypothesis-independent, can be absorbed into *a*(*t*) and *d* (*t*), correspondingly, and thus will not be considered further. As a result of this, the *y_i_* (*t*) used from now on is a “simplified” version of the log-likelihood.

We now take the logarithm of Eq 5, define – log*P_i_* (*t*) ≡ – log*P* (*H_i_*|x (1: *t*)) and substitute the simplified log-likelihood from Eq 8 in the result, giving:

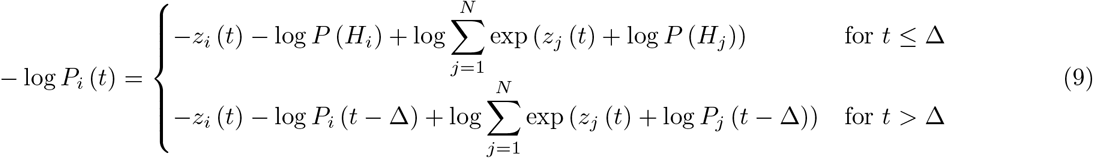

Where *z_i_* (*t*) = *y_i_* (*t*)+*c* (*t*) and the term *c*(*t*) models a hypothesis-independent baseline. Because of its uniformity across all hypotheses, *c*(*t*) has no effect on inference. It is defined in detail below.

The rMSPRT itself takes the form:

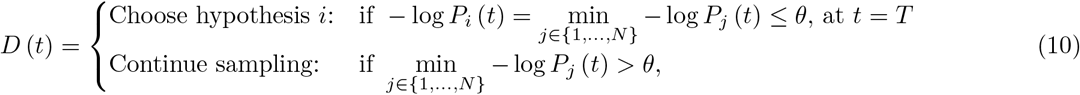

where *D*(*t*) is the decision at the discretely distributed time *t*, *θ* ∈ (0, – log (1/*N*)] is a constant threshold, and *T* is the decision termination time. Alternatively, an individual threshold per hypothesis can be set as {*θ*_1_,…, *θ*_*N*_}, giving a more general formulation.

### Cortical and thalamic baselines

According to our mapping of rMSPRT to the cortico-subcortical loops (Fig 5), the sensorimotor cortex baseline, *c*(*t*) (Eq 9), delayed with respect of the output of the model basal ganglia, is:

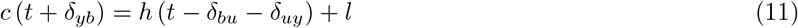

It houses a constant baseline l and the thalamo-cortical contribution *h*(*t* – *δ*_*bu*_ – δ_*uy*_), which in turn is the delayed cortical input to the thalamus

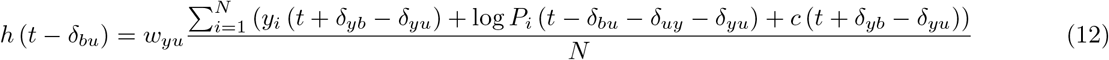

Here we have chosen *h*(*t* – δ_*bu*_) to be a scaled average of cortical contributions; nevertheless, any other hypothesis-independent function of them can be picked instead if necessary. It would thus not affect inference and render similar results.

The definitions above introduce two free parameters *l* ∈ℝ^+^ and *w_yu_* ∈ [0,1) that have the purpose of shaping the dynamics of the computations within rMSPRT during decision formation. The range of *w_yu_* ensures that the value of computations in the cortico-thalamo-cortical, positive-feedback loop is not amplified to the point of disrupting inference in the overall loop. Crucially, since both parameters are hypothesis-independent, none affects inference.

### Simulating the random dot motion task using the rMSPRT

For rMSPRT decisions to be comparable to those of monkeys, they must exhibit the same error rate, *ε* ∈ [0,1]. Error rates are taken to be an exponential function of coherence (%), s, fitted by non-linear least squares (*R^2^* > 0.99) to the behavioural psychometric curves from the analysed LIP database, including 0, 9, and 72.4 % coherence for this purpose. This resulted in:

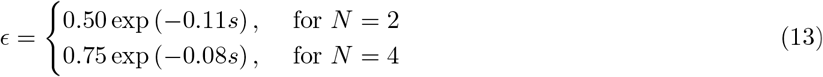

Since monkeys are trained to be unbiased regarding choosing either target, initial priors for rMSPRT are set flat (*P* (*H_i_*) = 1/*N* for all i) in every simulation. During each Monte Carlo experiment, rMSPRT made decisions with error rates from Eq 13. The value of the threshold, *θ*, was iteratively found to satisfy ∈ per condition. Decisions were made over data, *x_j_*(*t*)/*n*, randomly sampled from lognormal distributions specified for all channels by means and standard deviations *μ*_0_ and *σ*_0_, respectively; the exception was a single channel where the sampled distribution was specified by *μ*_*_ and *σ*_*_. This models the fact that MT neurons respond more vigorously to visual motion in their preferred direction compared to motion in a null direction, *e.g.* against the preferred. As explained in Fig 9, this effectively simulates macaque MT neural activity during the random dot motion task. The same parameters were used to specify likelihood functions per experiment.

#### Model parameters

To parameterize the input stochastic processes and likelihood functions of rMSPRT, we estimated the means μ_*_*n* and μ_0_*n*, and standard deviations *σ*_*_*n* and *σ*_0_*n* directly as those over the activity between 900 and 1900 ms after stimulus onset in the MT population, per condition (shown smoothed in Fig 1d,e). The subscript * indicates the condition when dots were predominantly moving in the direction preferred by the neuron. The subscript _0_ indicates when they were moving against it. We dub this parameter set Ω, and report it in Table 1. Fig 1f shows the lognormal ISI distributions specified by Ω; solid ones are *f*_*_, in our notation, and dashed ones are *f*_0_, per coherence.

We use *l* = 15, *w_yu_* = 0.4, *n* = 40, and *δ_yb_*, δ_*yu*_, δ_*bu*_, δ_*uy*_ = 1 (hence Δ = 3) in all simulations, unless otherwise stated. The value of latencies was set to 1 for simplicity. The values of the first three free parameters come from a manual tuning exercise with the aim of revealing a pattern in the model LIP akin to the ramp-and-fork one in LIP recordings; note that such a two-segment pattern is already guaranteed by the two cases of Eq 5.

The statistics in all simulations were from either the Ω or Ω_*d*_ parameter sets as noted per case. Note that the statistics actually used are those extracted from MT in Table 1, divided by the scaling factor *n*.

Second column: number of neurons for which data was available per coherence. *μ*: mean. *σ:* standard deviation. *n*: data scaling factor. Statistics with subscript * denote that dots were moving towards the preferred motion direction of the MT neuron, whereas _0_ denotes that they were moving in the opposite, null direction. The parameter set, iΩ (computed here from MT data) or Ω_*d*_ (after information depletion), to which each value corresponds is noted above them. Note that, due to the information depletion required to produce Ω_*d*_,μ_0_*n* and *σ*_0_*n* take different values for *N* = 2, 4.

### Spikes and continuous time interpretation of discrete time

We have defined rMSPRT to operate over a discrete time line; however, the brain operates over continuous time. [20] introduced a continuous-time generalization of MSPRT that uses spike-trains as inputs for decision. Thence, the length of ISIs is random and their sum up until decision is, by definition, a continuously distributed time. With all other assumptions equal, they demonstrated that, as an average, the traditional discrete-time MSPRT requires about the same number of observations to decision (discretely distributed), as the maximum number of ISIs among input channels, required by the more general spike-based MSPRT (also discretely distributed yet occurring over continuous time; Fig 9). This has two key implications. First, that continuous-time spike-trains can be substituted as decision evidence for (r)MSPRT by discrete-time stochastic processes —like **x**(*r*: *t*) here— as long as their distributions and the statistics that specify them remain equal; with this we gain efficiency on the implementation of discrete-versus continuous-time algorithms in digital computers, as well as simplicity on their analysis and interpretation. Second, and most important to compare the rMSPRT’s performance to experimentally-measured behaviour, that the (discretely distributed) number of observations to decision, *T*, in (r)MSPRT has an interpretation as continuously-distributed time. In brief, simulating decision evidence in discrete-time for (r)MSPRT as defined here is a simpler, equivalent way to simulate decisions made on the basis of continuous-time spike-trains. In light of this, the expected decision sample size for correct choices, 〈*T*〉_c_, required by the (r)MSPRT, can be interpreted as the mean decision time

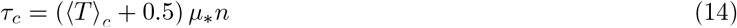

predicted by the more general continuous-time, spike-based MSPRT, where μ_*_*n* is the mean ISI produced by a MT neuron whose preferred motion direction was matched by the stimulus and was thus firing the fastest on average (Fig 9). When the mean firing rate to a preferred characteristic of the stimulus is larger than that to a non-preferred one (μ_*_ < μ_0_) —as in MT [24], middle-lateral, and anterolateral auditory cortex [66]— the hypothesis selected in error trials is that misinformed by channels with mean μ_0_*n* which intuitively happened to fire faster than those whose mean was actually μ_*_*n*. Hence, the mean decision time predicted by rMSPRT in error trials would be:

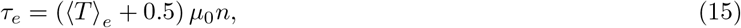

where 〈T〉 is the mean decision sample size for error trials. An instance of rMSPRT capable of making choices upon sequences of spike-trains is straightforward from the formal framework above and that introduced by [20]; nevertheless, as said, for simplicity here we choose to work with the discrete-time rMSPRT. After all, thanks to Eqs 14 and 15 we can still interpret its behaviour-relevant predictions in terms of continuous time. These are used to compute decision times throughout.

### Estimation of lost information

We outline here how we use the monkeys’ reaction times on correct trials and the properties of the rMSPRT, to estimate the amount of discrimination information lost by the animals. That is, the gap between all the information available in the responses of MT neurons, as fully used by the rMSPRT (parameter set Ω), and the fraction of such information actually used by monkeys.

The expected number of observations to reach a correct decision for (r)MSPRT, 〈T〉_*c*_, depends on two quantities. First, the mean total discrimination information required for the decision, *I* (∈, *N*), that depends only on the error rate, ∈, and *N*. Second, the ‘distance’ between distributions of ISIs from MT neurons that are simultaneously contributing evidence for decision, while visual motion matches the tuning of some and not others (*e.g.* red versus black in Fig 9). This distance is the Kullback-Leibler divergence from *f*_*_ to *f*_0_

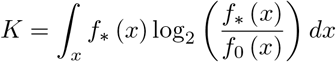

which measures the discrimination information available between the distributions. Using these two quantities, the decision time in the (r)MSPRT is [20]:

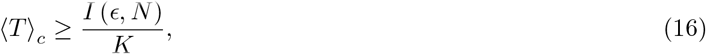

The product of our Monte Carlo estimate of 〈T〉 in the rMSPRT (Fig 3a in the Results) and *K* from the MT ISI distributions (Fig 1f), gives an estimate of the limit *I* (∈, *N*) in expression 16, denoted by 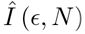.

The ‘mean decision sample size’ of monkeys —hence the superscript ^m^— within this framework corresponds to 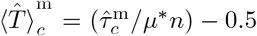 (from Eq 14). Here, 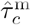 is the estimate of the mean decision time of monkeys for correct choices, per condition; that is, the reaction time from Fig 3a minus some constant non-decision time. With 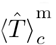 and 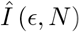, we can estimate the corresponding discrimination information available to the monkeys in this framework as 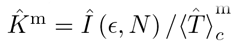 (from expression 16).

Fig 3b compares K (red line) to 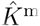 (blue/green lines and shadings) for monkeys, using non-decision times in a plausible range of 200-300 ms. Fig 3c shows the discrimination information lost by monkeys as the percentage of 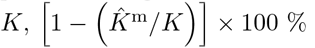

### Information depletion procedure

Expression 16 implies that the reaction times predicted by rMSPRT should match those of monkeys if we make the algorithm lose as much information as the monkeys did. We did this by producing a new parameter set that brings *f*_0_ closer to *f*_*_ per condition, assuming 250 ms of non-decision time; critically, simulations like those in Fig 4 will give about the same rMSPRT reaction times regardless of the non-decision time chosen, as long as it is the same assumed in the estimation of lost information and this information-depletion procedure.

An example of the results of information depletion in one condition is in Fig 3d. We start with the original parameter set extracted from MT recordings, Ω ( ‘preferred’ and ‘null’ densities in Fig 3d), and keep μ_*_ and σ_*_ fixed. Then, we iteratively reduce or increase the differences |μ_0_ – μ_*_| and |σ_0_ – σ_*_| by the same proportion, until we get new parameters μ_0_ and σ_0_ that, together with μ_*_and σ_*_, specify preferred ( ‘preferred’ in Fig 3d) and null ( ‘new null’) density functions that bear the same discrimination information estimated for monkeys, 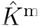; hence, they exactly match the information loss in the solid lines in Fig 3c. Intuitively, since the ‘new null’ distribution in Fig 3d is more similar to the ‘preferred’ one than the ‘null’, the Kullback-Leibler divergence between the first two is smaller than that between the latter two. The resulting parameter set is dubbed Ω_*d*_ and reported in Table 1. Note that this is not a fitting procedure, which would be merely descriptive. Instead, information depletion takes advantage of the properties of the (r)MSPRT to describe the data, but also to predict that the longer decision times of monkeys are explained by a reduction in the discrimination information in the streams of decision evidence.

### Information loss for an enhanced match of reaction times

The slight deviation of the mean reaction times of (r)MSPRT vs those of monkeys in Fig 4a stems from the expression 16 being an inequality. Due to this, 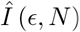 is a likely over-estimate of *I* (∈, *N*). Dividing 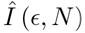 by 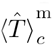 hence gives an over-estimate of the monkey discrimination information, 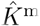. If then rMSPRT uses statistics consistent with this over-estimated 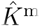, it renders under-estimated reaction times. This residual discrepancy can be corrected by further multiplying 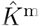, per condition, by the corresponding ratio of the decision time of the model over that of the monkey, from Fig 4a. Repeating the simulations with the implied parameter set would trivially render rMSPRT reaction times that more exactly match those of monkeys (not shown). This will likely carry with it a better match in error trials, which is unconstrained in the procedure. Nonetheless, this exercise gives us the information loss associated to such enhanced match, shown in Fig 3c as dashed lines for a 250 ms non-decision time (compare to solid lines); this constitutes a further refined measure of the minimum information lost by the animals according to our framework.

## Supplementary Information

S1 Fig. Mean reaction times in error trials are consistently longer than those of correct trials (Fig 10).

**Figure 10:**
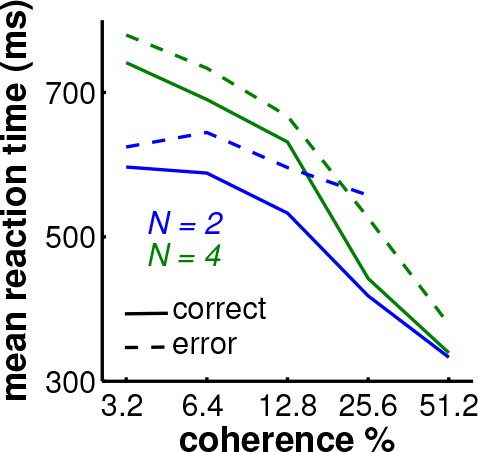
Mean error reaction times are longer than correct ones. Monkey mean reaction times from Fig 4, panels a and b, shown in the same plot for comparison. Solid: correct trials. Dashed: error trials. Blue, green: *N* = 2, 4.

S2 Fig. Predicted contribution of the corticothalamic projection (Fig 11).

**Figure 11:**
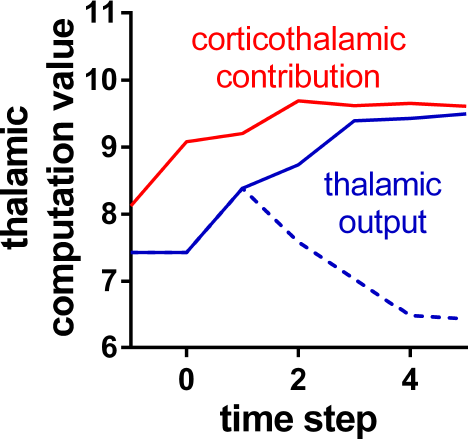
Influence of cortico-thalamic contribution in rMSPRT. Example mean cortico-thalamic contribution, *h*(*t* – δ;_*bu*_) (red), compared to the mean thalamic output during inRF settings (solid blue) and outRF ones (dashed blue) for 25 % coherence and *N* = 2. Single Monte Carlo experiment with 800 total trials.

S3 **Fig. Mapping of rMSPRT** computations to basal ganglia nuclei (Fig 12).

**Figure 12:**
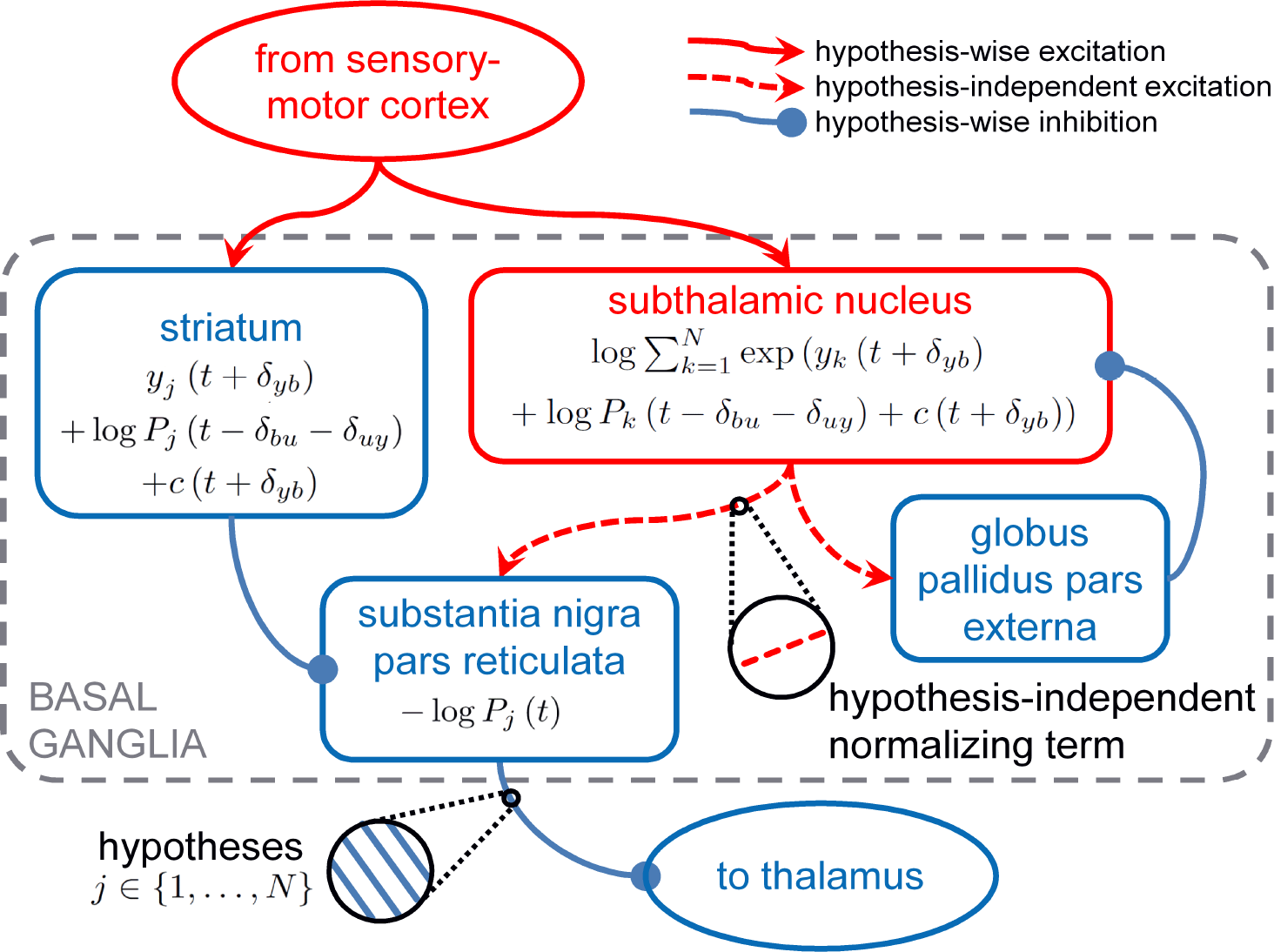
Mapping of rMSPRT computations to the basal ganglia. Parallel computations for N hypotheses —indexed by *j*— mapped onto the basal ganglia nuclei, within the grey dashed box (see [17,18,62]). This may include the pathway from striatum to the globus pallidus pars externa, as well as the pathway from the latter to substantia nigra pars reticulata, as demonstrated by [17], thus representing all major pathways among the basal ganglia. It has recently been shown that neurons in the microcircuitry joining the rodent subthalamic nucleus and globus pallidus, are theoretically able to collectively represent the normalization term required by algorithms in the family of the (r)MSPRT [21]; this, as well showing that the mapping of MSPRT (thus rMSPRT) to the basal ganglia can also account for a connection from globus pallidus pars externa to striatum. Same conventions and notation as in Fig. 5. All computations are delayed with respect to the substantia nigra pars reticulata. 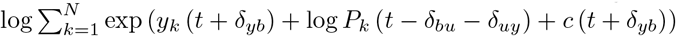: normalization term (from Eq 9), putting together the cortical computations for all hypotheses into a hypothesis-independent contribution. Note that the model striatum represents a copy of the cortical signal (as in Fig. 5) but its influence on the substantia nigra pars reticulata is the negative of such cortical input.

S4 Fig. Neural populations in MT are reliably modulated by coherence in the random dot motion task as soon as 50 ms after stimulus onset (Fig 13).

**Figure 13:**
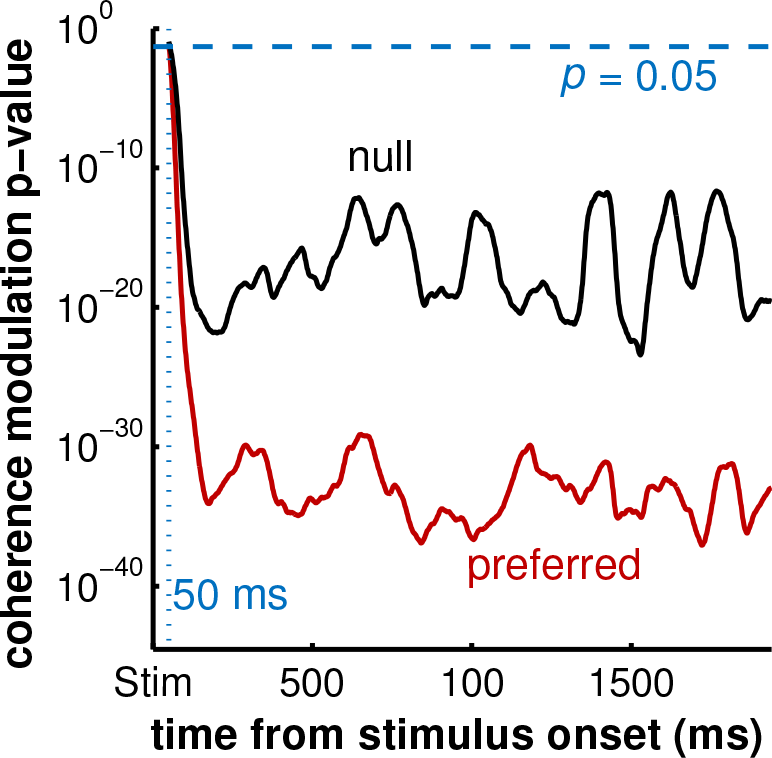
MT neurons are significantly modulated by coherence as soon as 50 ms after dots appear in the random dot motion task. With the method described in the main article, we computed the mean firing rate of every MT neuron in our data facing dots moving in its preferred or null motion directions at six coherence levels (%): 0, 3.2, 6.4, 12.8, 25.6, 51.2 (randomly assigning 0% trials between directions). For every neuron, every 1 ms bin, we conducted a linear regression per motion direction of the form *v* = *β* + *β*_s_s, where *v* are the mean firing rates at every bin, *s* are the corresponding coherence levels (%), and *β*, *β*_s_ are the intercept and the coefficient for the coherence contribution, respectively. We then applied a *t*–test where the null hypothesis was: the mean of the distribution of the 189–213 *β*_s_’s (again, one per MT neuron) we got per direction, equals 0. Here we show the corresponding *p*–value for MT coherence modulation, per direction, aligned at stimulus onset (Stim). Note that this involved conducting a single statistical test/comparison for every 1 ms bin, independent of those conducted in surrounding bins. Red: dots moving in the preferred direction of recorded neuron. Black: moving in the opposite, null direction. Horizontal blue dashed line: *p* = 0.05. Vertical blue dotted line: 50 ms. The coherence modulation p-values for preferred and null directions drop under 0.05 about ∼ 50 ms after the onset of the dots stimulus and drop much further soon after this.

## Acknowledgments

We thank Anne Churchland, Roozbeh Kiani, Michael Shadlen, Long Ding, and Joshua Gold for sharing their experimental data and the Humphries lab (Abhinav Singh, Mathew Evans, and Silvia Maggi), Rafal Bogacz, and Long Ding for discussions. This work was supported by a National Council of Science and Technology (CONA-CyT) Fellowship (JAC), a Medical Research Council Senior non-Clinical Fellowship (MDH; MR/J008648/1), and a Wellcome Trust collaborative research award (JAC, MDH; 105610/Z/14/Z).

